# Fluorinated cGAMP analogs, which act as STING agonists and are not cleavable by poxins: structural basis of their function

**DOI:** 10.1101/2023.09.07.556653

**Authors:** Martin Klima, Milan Dejmek, Vojtech Duchoslav, Andrea Eisenreichova, Michal Sala, Karel Chalupsky, Dominika Chalupska, Barbora Novotná, Gabriel Birkuš, Radim Nencka, Evzen Boura

**Affiliations:** Institute of Organic Chemistry and Biochemistry AS CR, v.v.i., Flemingovo nam. 2., 166 10 Prague 6, Czech Republic

## Abstract

The Stimulator of Interferon Genes (STING) plays a crucial role in the cGAS-STING pathway of innate immunity, detecting DNA in the cytoplasm and defending against certain cancers, viruses, and bacteria. We designed and synthesized fluorinated carbocyclic cGAMP analogs, MD1203 and MD1202D (MDs), to enhance their stability against nucleases and their affinity for STING. These compounds demonstrated exceptional activity against wild-type STING and all its allelic variations, including the hard-to-target REF isoform. Despite their distinct chemical modifications relative to the canonical CDNs, such as the substitution of guanine with hypoxanthine and the fluorination of the (pseudo)ribose ring, crystallographic analysis revealed a consistent binding mode with STING. Importantly, these compounds were resistant to cleavage by viral poxin nucleases. The crystallographic analysis of poxin/MD complexes unveiled their binding mode at the interface of poxin monomers, with dynamic adenine base orientations. Interestingly, MDs-bound poxin adopted an unliganded-like conformation, distinct from the conformation of cGAMP-bound poxin. Moreover, when MDs were in complex with poxin, they exhibited a different conformation than cGAMP when bound to poxin; in fact, it closely resembled the conformation observed when MDs were bound to STING. In conclusion, the development of MD1203 and MD1202D, showcases their potential as potent STING activators with remarkable stability against poxin-mediated degradation—a crucial characteristic for future development of antivirals.

## Introduction

Viruses pose a formidable threat to organisms across the evolutionary spectrum, including humans. They have evolved intricate strategies to infiltrate cells in order to replicate. Consequently, cells must possess a robust defense arsenal. One pivotal defense mechanism is the production of type I interferons (IFN I), which play a fundamental role in establishing an antiviral response, including alerting neighboring cells to the presence of viral infection [1, 2].

Central to the induction of interferon production lies the cGAS-STING pathway, an essential molecular cascade in which cGAS (cyclic GMP-AMP synthase) serves as an innate immune sensor, adept at recognizing cytosolic dsDNA [3, 4]. This includes pathogen-derived DNA arising from bacteria, DNA viruses, or reverse transcription of retroviruses, as well as self-DNA emanating from damaged or deceased cell nuclei or mitochondria [5]. Upon detection of cytosolic DNA, cGAS undergoes dimerization and generates 2’,3’-cyclic-GMP-AMP (cGAMP), which is a second messenger [6, 7]. Once cGAMP binds to STING (stimulator of interferon genes), it initiates the entire signaling cascade. STING translocates from the endoplasmic reticulum to the Golgi, where it recruits and activates the TBK1 kinase. Consequently, TBK1 phosphorylates IRF3, culminating in its dimerization and nuclear translocation [8]. Once within the nucleus, IRF3 binds to specific DNA sequences, thereby initiating the transcriptional activation of type I interferon genes [9].

However, viruses, in the course of their evolutionary trajectory, have developed strategies to overcome and undermine these intrinsic immune defenses of their host organisms [3].

Poxins, a family of unusual nucleases that are specific towards cGAMP, have been recently discovered in in pox viruses (hence the name) [10]. These enzymes enable the viruses to evade the host’s innate immune system by degrading cGAMP and leading to the inhibition of the cGAS-STING pathway [11]. In detail, crystal structures of vaccinia virus poxin in pre- and post-reactive states define the mechanism of selective cGAMP degradation through metal-independent cleavage of the 3′-5′ bond, converting cGAMP into linear Gp[2′-5′]Ap[3′] [10]. Thus, poxins are used by viruses as an evasion mechanism which allows the viruses to dampen the immune response mediated by cGAMP and potentially establish successful infection.

Recently, the Carell team observed that dideoxy-2′,3′-cGAMP and its derivatives, characterized by the absence of secondary ribose-OH groups, constitute a class of STING agonists that demonstrate stability against poxins [12]. We have prepared a novel class of fluorinated cyclic dinucleotides that exhibit high potency against STING and are not cleavable by poxin. We have solved crystal structures of these compounds in complex with both STING and monkeypox virus (MpxV) poxin that explain their potency and stability.

## Results

### Fluorinated cGAMP analogs

We decided to prepare carbocyclic cGAMP analogs fluorinated at the (pseudo)sugar rings because both of these modifications could lead to the increase in stability of the molecules [13] as well as in the affinity towards STING [14]. Through a straightforward, yet rather challenging syntheses (**Scheme S1**) starting from a protected Vince Lactam **1** we prepared the key intermediate **10** (**Fig. 1**). This monomer was coupled and subsequently cyclized with 2′-fluoro-2′-deoxyadenosine phosphoramidite. The standard means of connecting both (thio)phosphate bonds were employed. The linear dinucleotide was formed using py-TFA/tBHP and py-TFA/xanthane hydride systems, respectively. Subsequently, the macrocycle was closed with DMOCP/iodine and DMOCP/Beaucage reagent sequence, respectively (**Fig. 1, Scheme S2**). This resulted in the cyclic dinucleotides MD1203 and MD1202D (further referred to also as MDs) after deprotection using methylamine.

**Figure 1.**
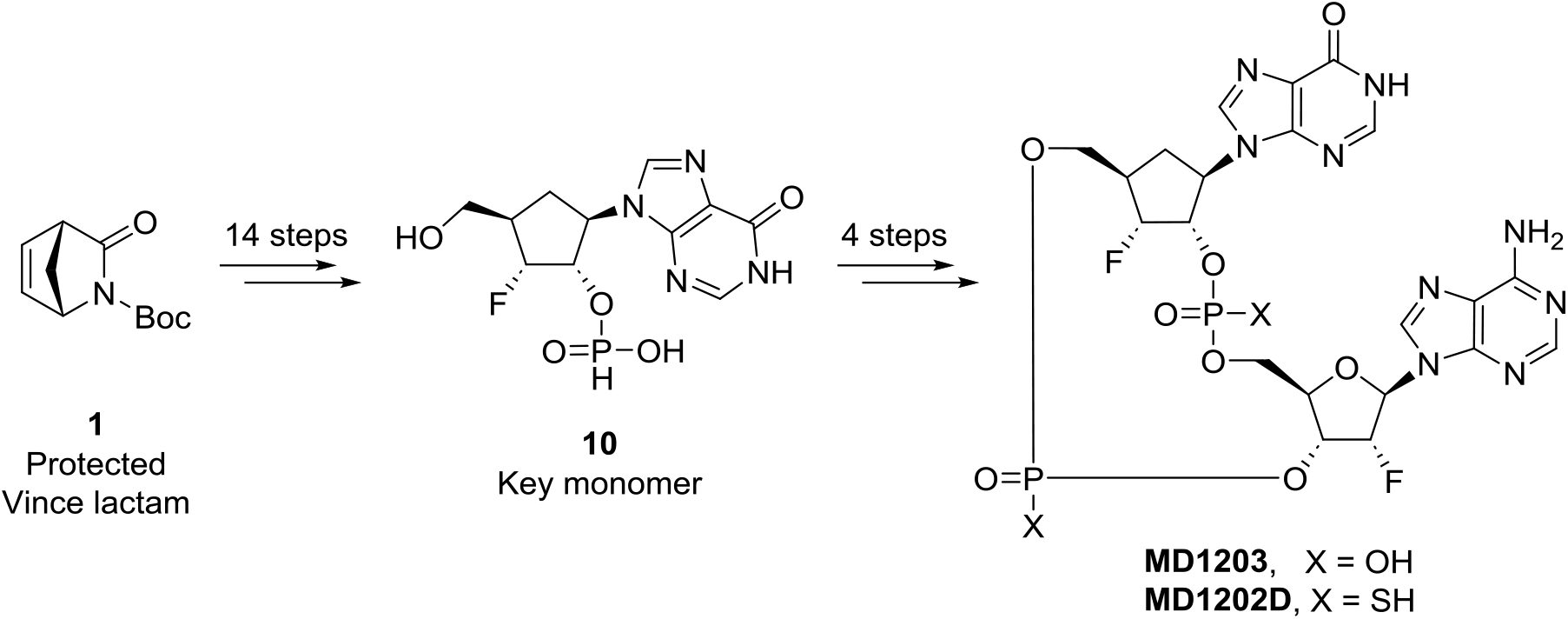
Simplified scheme of synthesis of MD1203 and MD1202D. The overall synthesis used a protected Vince lactam as starting material to synthesize a monomer that was subsequently chemically cyclized. Details are in **Schemes S1, S2**.

### STING activity assay

The agonistic activity towards wild-type STING and its most abundant allelic forms (HAQ, REF, AQ, Q) was evaluated using a cell-based reporter assay, in which we included digitonin A, a mild detergent, for cell membrane permeabilization [15-17] (**Table 1**). Without this permeabilization the negatively charged CDNs enter the cells only slowly, which significantly decreases their effectiveness. Both MD1203 and MD1202D proved to be very strong activators of STING across the spectrum of allelic variants, with their EC_50_ values similar or better than canonical CDNs and the former clinical candidate ADU-S100 (Table 1).

**Table 1.** *In vitro* activity of selected CDNs in the digitonin HEK 293T cell-based reporter assay.

|  | Base | R | X | WT<br>(μM) | HAQ<br>(μM) | REF<br>(μM) | AQ<br>(μM) | Q (μM) |
| --- | --- | --- | --- | --- | --- | --- | --- | --- |
| 2'3'cGAMP | G | OH | O | 0.02<br>(±0.004) | 0.02<br>(±0.003) | 0.07<br>(±0.02) | 0.04<br>(±0.01) | 0.05<br>(±0.02) |
| ADU-S100 | A | OH | S | 0.08<br>(±0.01) | 0.26<br>(±0.05) | 1.64<br>(±0.28) | 0.23<br>(±0.03) | 1.01<br>(±0.19) |
| 2'3'-c-dGdAMP | G | H | O | 0.05<br>(±0.018) | 0.02<br>(±0.011) | 3.3<br>(±0.9) | 0.03<br>(±0.01) | 0.06<br>(±0.03) |
| MD1202D | Hx | F | S | 0.02<br>(±0.009) | 0.03<br>(±0.014) | 0.36<br>(±0.17) | 0.02<br>(±0.007) | 0.38<br>(±0.22) |
| MD1203 | Hx | F | O | 0.03<br>(±0.016) | 0.06<br>(±0.03) | 2.45<br>(±0.81) | 0.06<br>(±0.02) | 0.85<br>(±0.55) |

<sup>a</sup>Results from the digitonin assay using 293T reporter cells expressing different STING protein haplotypes; EC<sub>50</sub> values are expressed as mean ± SEM of at least two independent experiments measured in triplicates.

### Structural basis for the activation of STING by MD1203 and MD1202D

The atomic details of the interaction of STING with MDs were uncovered by the crystallographic analysis of the respective protein-ligand complexes. The obtained structures were in a good agreement with previous crystallographic studies on the STING protein complexed with CDNs. MD1203 and MD1202D were bound to STING in a usual stoichiometry protein:CDN 2:1 [18]. The dimer of the STING CDN-binding domain was present in the closed conformation with the lid formed above the ligand-binding site (**Fig. 2a**). The binding of the adenine and guanine bases of MD1203 and MD1202D was mediated mainly by parallel π-π stacking with Tyr167 and by π-cation interactions and hydrogen bonds with Arg238. The macrocycles of MD1203 and MD1202D interacted with STING via hydrogen bonds and salt bridges with Ser162, Thr263, Thr267, and Arg238 (**Fig. 2b-c**). Overall, the atomic details of the interactions of between STING and MD1203, as well as MD1202D, exhibited similarities to the previously published structures of the STING-CDN complexes [14, 17, 19, 20].

**Figure 2.**
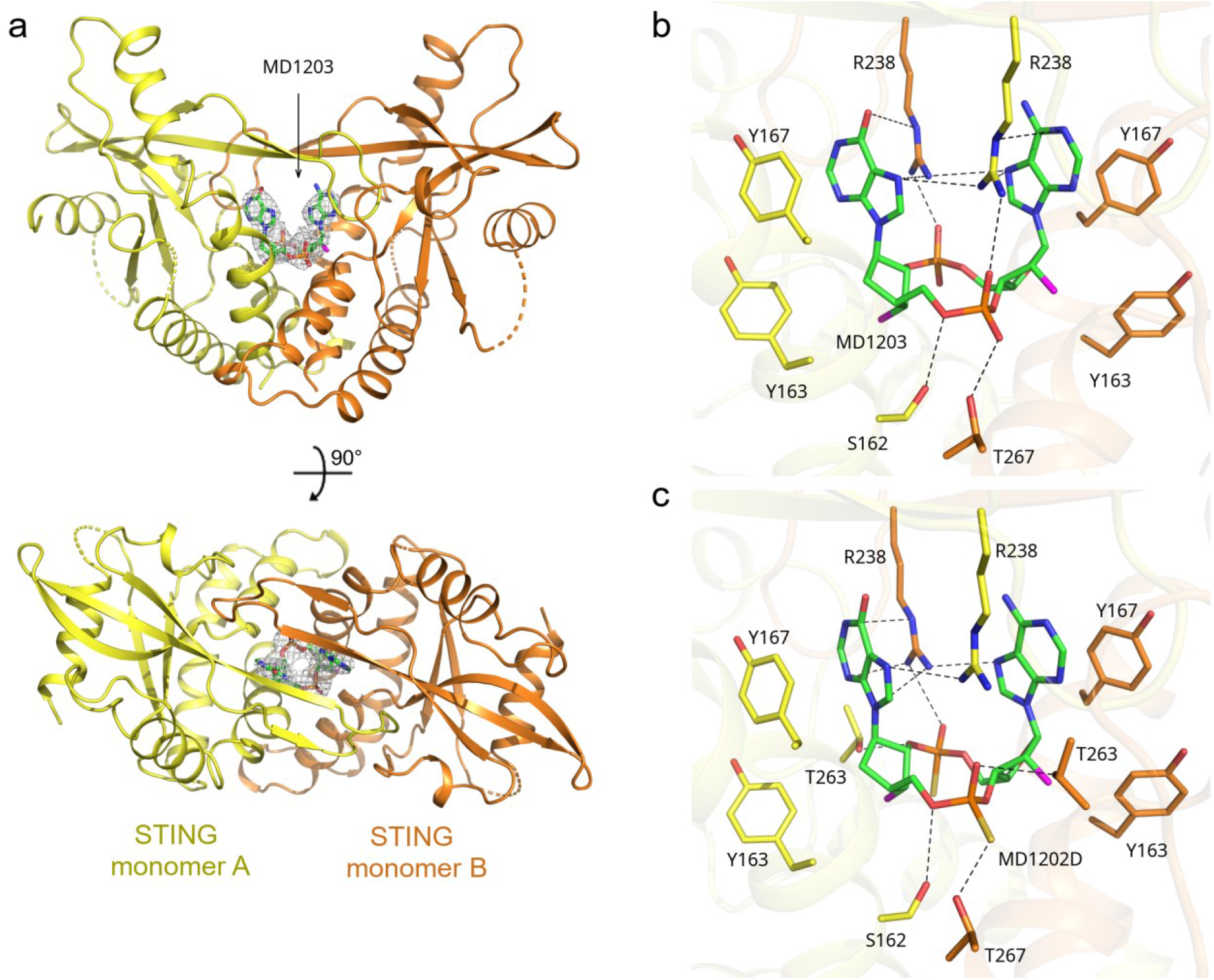
Crystal structure of human STING in complex with MD1203 and MD1202D. **a**) Overall view of the STING/MD1203 complex. The protein backbone is shown in cartoon representation; the two STING monomers are depicted in yellow and orange. MD1203 is shown in stick representation and colored according to elements: carbon, green; nitrogen, blue; oxygen, red; phosphorus, orange; fluorine; magenta. The *Fo*-*Fc* omit map contoured at 3σ is shown around the ligand. **b-c**) Detailed view of the ligand binding site. MD1203 (**b**) or MD1202D (**c**) and side chains of selected STING residues are shown in stick representation with carbon atoms colored according to the protein/ligand assignment and other elements colored as in **a**. Selected hydrogen bonds involved in the STING-ligand interaction are presented as dashed black lines.

### Enzymatic stability of MD1203 and MD1202D

As mentioned above, viruses use special nucleases, poxins, to degrade cGAMP. However, we speculated that the unique features of MDs, in which the (pseudo)sugar hydroxyl groups were replaced by fluorine atoms, will make these CDNs resistant to poxin-mediated hydrolysis, while retaining the STING agonistic activity. The highly electronegative fluorine atom serves as a bioisosteric substitute for the hydroxyl group, closely mimicking the naturally occurring CDNs [14]. We prepared a recombinant MpxV poxin and directly tested the stability of MDs in its presence. While cGAMP was rapidly degraded by the poxin nuclease, both MD1203 and MD1202D exhibited remarkable stability, with no degradation observed after one hour of incubation (**Fig. S1**).

### MD1203 and MD1202D recognition by MpxV poxin

These compounds are not cleaved by poxins. However, we speculated that due to their chemical structure, which inherently resembles cGAMP, they would still bind and be recognized by poxins. Crystallography is increasingly used as an analytical tool by us and others [21-24]. We attempted to obtain crystals of MD1203 and MD1202D complexed with poxins to gain molecular insights into the poxins’ recognition mechanism of these CDNs. Indeed, we successfully obtained well-diffracting crystals (1.65Å and 1.93Å, respectively, **Table S1**) for these complexes. The structures were determined using molecular replacement (detailed in the Materials and Methods section) and were refined to favorable R-factors and geometry (**Table S1**).

Poxins exist as dimers, and the binding site for CDNs is situated at the monomer-monomer interface [10, 11]. As expected, both MDs were located at this interface and their density was clearly visible (**Fig. 3**). Both MDs, like cGAMP, are non-symmetric; they have a hypoxanthine and an adenine base. The binding mode of the hypoxanthine base is the same in both structures (**Fig. 3b, c**), its oxygen atoms forms a hydrogen bond with the sidechain of His17 and also binds the nitrogen atom of Ala18 (in the case of MD1203 through a water bridge or, in the case of MD1202D, directly), another water bridge is formed to Tyr138 of the other poxin monomer. This monomer also forms several water bridges, mainly through Arg182, Arg184, Gln169 and Gly127, to the macrocycles of MD1202D and MD1203. However, the adenine base adapts a different orientation in each structure. It undergoes an approximate 180-degree rotation relative to the adenine of the other CDN. Intriguingly, both conformations facilitate π-π stacking with the aromatic ring of Tyr138. But only the conformation observed in the case of MD1202D allows also for a hydrogen bond to His17 (**Fig. 3c**). Nevertheless, both conformations are allowed and, most likely, exist in a dynamic equilibrium. This is supported by the fact that the B-factors for the adenine base are notably high (55 to 60), implying considerable mobility. In contrast, the B-factors for the hypoxanthine base are comparatively low (30 to 40), suggesting that this base is well fixed in its position (**Fig. 3d, e**).

**Figure 3.**
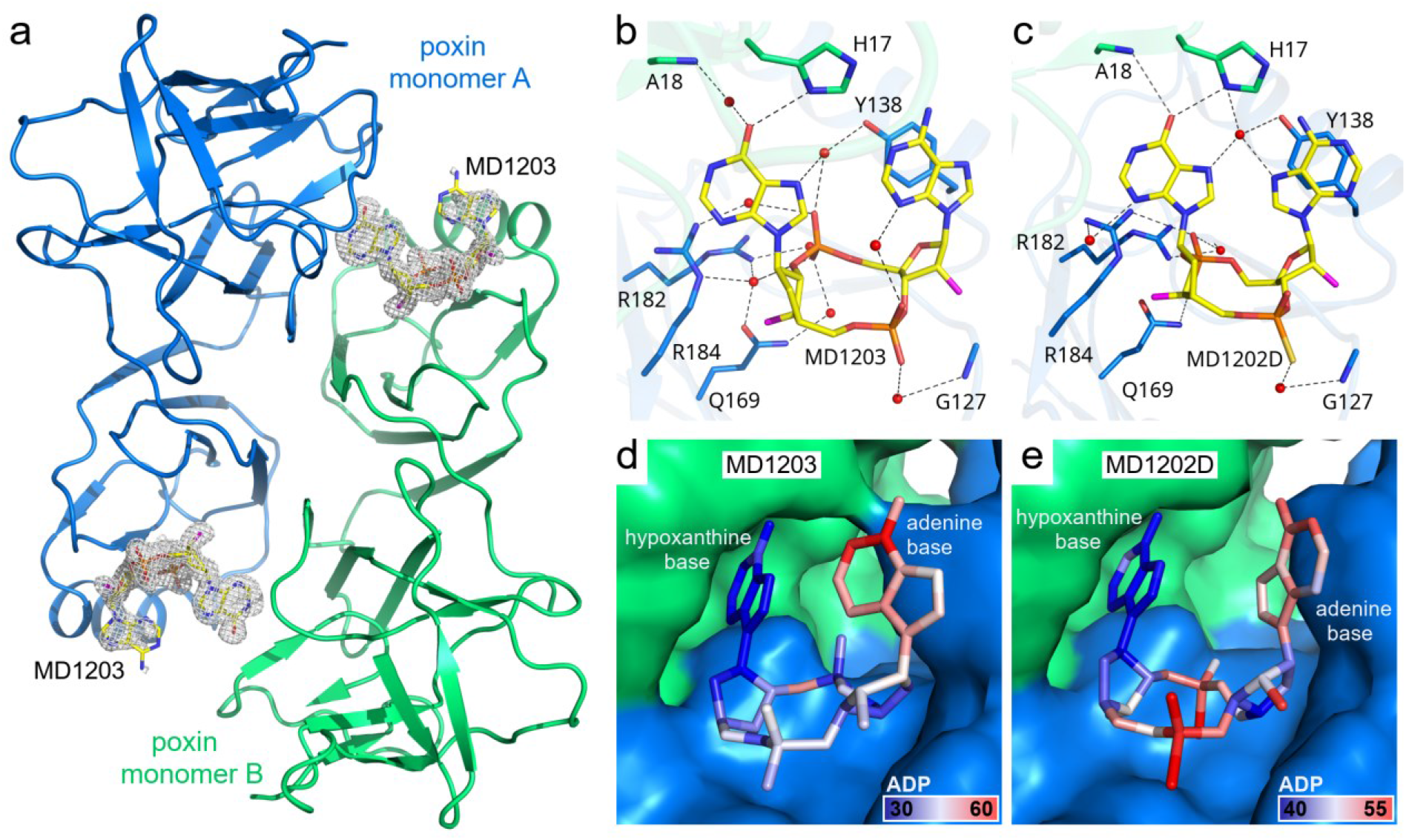
Crystal structure of MpxV poxin in complex with MD1203. **a)** Overall view of the poxin/MD1203 complex depicted as in **Fig. 2a**. The two poxin monomers are colored in blue and green, MD1203 carbons in yellow. The *Fo-Fc* omit map contoured at 3σ is shown around the two ligands. **b-c**) Detailed view of the MD1203 (**b**) and MD1202D (**c**) ligand binding site depicted as in **Fig. 2b. d-e**) MD1203 (**d**) or MD1202D (**e**) are colored according to the atomic displacement parameters (ADPs, also known as *B* factors) using the gradient from blue (atoms with lowest ADPs) to red (atoms with highest ADPs). The protein molecules are shown in surface representation.

We expected that MDs-bound poxin would adopt the same conformation as cGAMP-bound poxin. Interestingly, this was not the case. cGAMP binding induces a movement of the α1 helix (by approximately 3 Å), causing a narrowing of the CDN binding site (**Fig. S2**). However, upon structural comparison of MDs-bound or cGAMP-bound poxin with the unliganded poxin, we found that MDs-bound poxin adopts an unliganded-like conformation (**Fig. 4, Fig. S2**) and also the conformation of poxin bound MD compounds is different from that of poxin bound cGAMP (**Fig. 4c**). Structurally, in case of the cGAMP, the guanine base forms a hydrogen bond with Ala18 through its amine group and with Lys186 through its oxygen atom, while Lys186 is not involved in the recognition of the hypoxanthine base in MDs. The adenine base forms a hydrogen bond with Asn149 in the case of cGAMP. However, in the case of MDs, yet again, this residue is not involved in the recognition of the adenine base. On the contrary, Tyr138, which is involved in the macrocycle binding in the case of cGAMP, is observed to participate in π-π stacking interactions with the adenine base of MDs. Chemically, the major differences between MDs and cGAMP are the exchange of the guanine base for the hypoxanthine base and fluorination of the ribose rings. It is unlikely that the hypoxanthine-guanine exchange is responsible for the different binding mode as the amine group of the adenine base does not form any bond with poxin [10]. The carba modification of one of the ribose rings is also unlikely to have caused this change, as this structural feature has never significantly affected behavior of CDNs towards STING [25, 26]. However, fluorination of the (pseudo)ribose ring significantly improved binding of CDNs to STING by altering the enthalpy and entropy contributions. Interestingly, the crystal structures revealed the same binding conformation of STING [14], which we also observed for MDs binding towards STING (**Fig. 5**).

**Figure 4.**
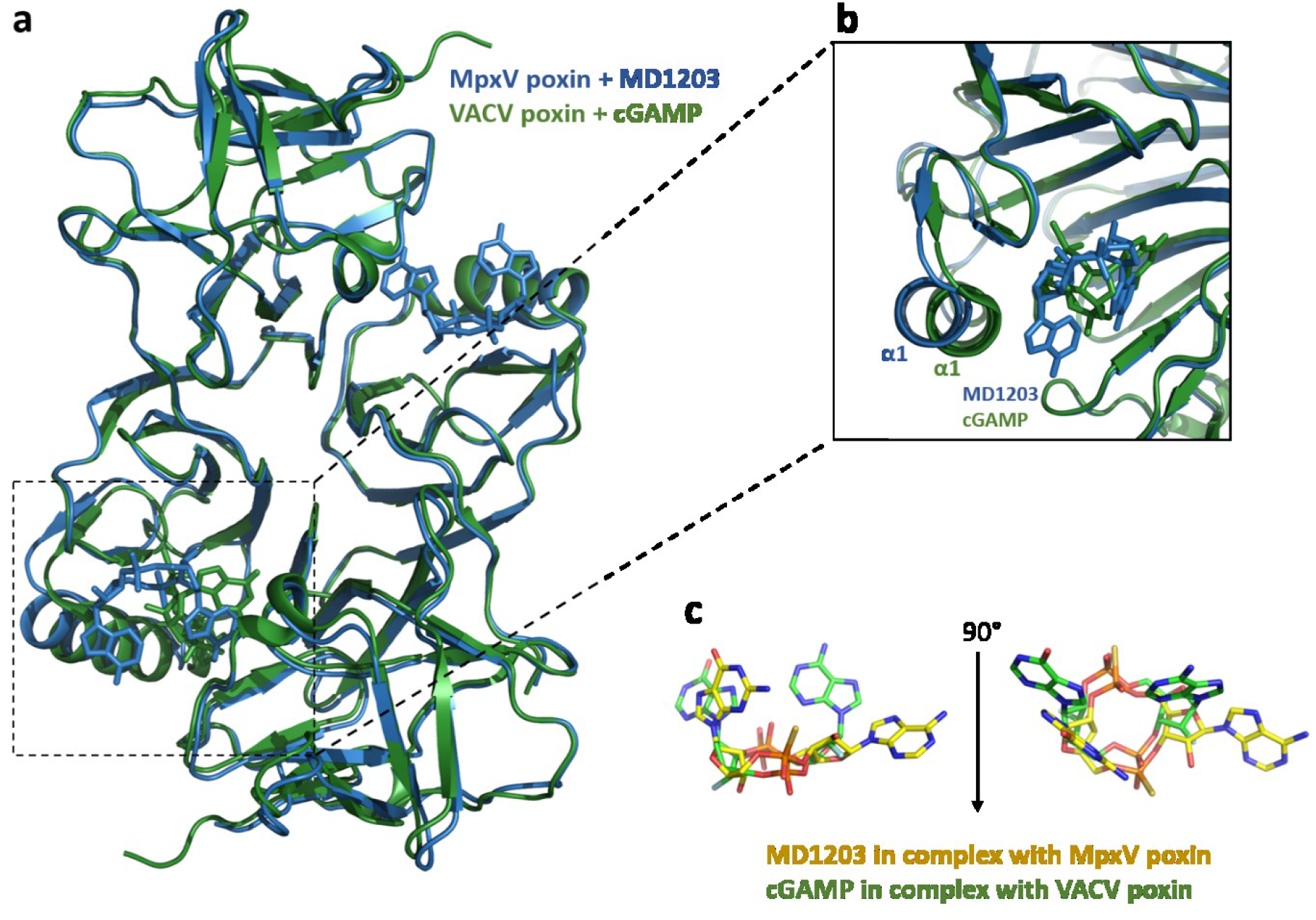
Conformational changes induced in VACV poxin by ligand binding. **a)** Structural alignment of VACV poxin in complex with cGAMP, depicted in green (PDB code: 6EA8), and MpxV poxin in complex with MD1203, depicted in blue. cGAMP binding to the VACV poxin induces conformational changes of α-helix 1.**b)** Close-up on the conformational changes of α-helix 1. **c)** Superposition of cGAMP bound to VACV poxin and MD1203 bound to MpxV poxin. cGAMP is depicted in green and MD1203 in yellow.

**Figure 5.**
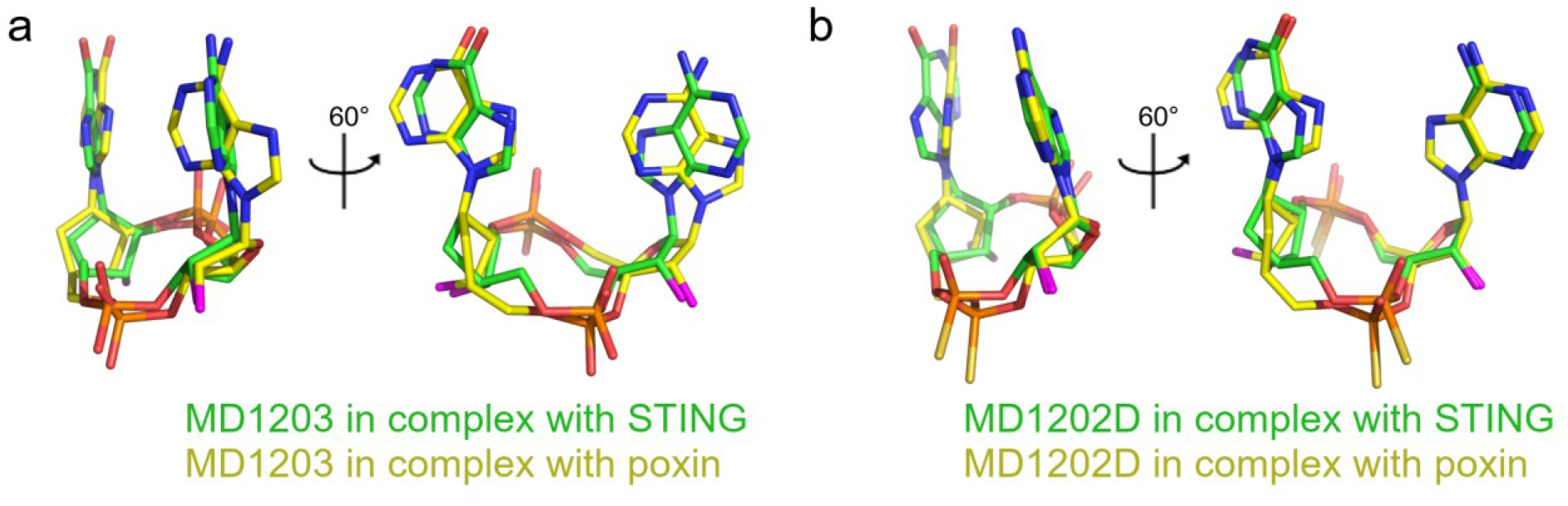
Conformations of MD1203 and MD1202D. **a)** Superposition of MD1203 bound to poxin and STING. MD1203 is shown in stick representation with carbon atoms colored in yellow (in complex with poxin) or green (in complex with STING) and other elements colored as in **Fig 2a**. The protein molecules are not shown. **b)** Superposition of MD1202D bound to poxin and STING depicted as in **a**.

## Materials and Methods

### Synthesis

Conventional techniques of organic chemistry were employed and are described in detail, along with the NMR spectra, in the Supplementary Information.

### Protein expression and purification

Poxin (amino acid residues 140-343) and STING (amino acid residues 1-197) proteins were purified using the same protocols as before [27, 28]. Briefly, they were expressed as fusion proteins with the 8xHis-SUMO tag, which facilitates purification and enhances solubility. The proteins were expressed in the *E. coli* BL21 DE3 Star and NiCo bacterial strains, respectively, in the autoinduction ZY-5052 medium. Bacterial cells expressing poxin or STING were harvested by centrifugation and disrupted by sonication or using the Emulsiflex C3 instrument (Avestin) in the lysis buffer (50 mM Tris pH 8.0, 300 mM NaCl, 20 mM imidazole, 10% glycerol, 3 mM β-mercaptoethanol), respectively. The lysates were precleared with centrifugation for 30 min at 30,000 g and incubated with the HisPur Ni-NTA Superflow agarose (Thermo Fisher Scientific) for 60 min. Then, the agarose beads were extensively washed with the lysis buffer (STING) or with the lysis buffer containing 1 M NaCl (poxin) and the proteins of interest were eluted with the elution buffer (50 mM Tris pH 8, 300 mM NaCl, 300 mM imidazole, 10% glycerol, 3 mM β-mercaptoethanol). The His_8_-SUMO tag was cleaved overnight with the recombinant yeast Ulp1 protease. Poxin was dialyzed against the lysis buffer and the 8xHis-SUMO tag was removed by reverse Ni-NTA chromatography. The resulting protein was further purified using the size exclusion chromatography at the HiLoad 16/600 Superdex 75 prep grade column (Cytiva) in the SEC buffer (20 mM HEPES pH 7.5, 250 mM KCl, 0.5 mM TCEP). STING was purified using the size exclusion chromatography at the HiLoad 16/600 Superdex 75 prep grade column (Cytiva) in the buffer A (50 mM Tris pH 7.4, 50 mM NaCl) followed by the anion exchange chromatography at the HiTrap Q HP column (Cytiva) using a gradient from the buffer A to the buffer B (50 mM Tris pH 7.4, 1M NaCl). The purified proteins were concentrated to 15-20 mg/ml, aliquoted, flash frozen in liquid nitrogen, and stored at 193 K until required.

### Crystallization and crystallographic analysis

Poxin was crystallized as before [27] except that it was supplemented with 2 mM MD1203 or MD1202D while STING was supplemented with 10 mM EDTA, which improves the quality of STING crystals [28], and 1 mM MD1203 or MD1202D. Diffraction quality crystals were obtained upon seeding in sitting drops that were prepared using 300 nl of protein with the indicated ligand and 300 nl of mother liquor (for poxin/MD1203: 100 mM phosphate-citrate pH 4.2, 40% PEG 300; for poxin/MD1202D: 200 mM NH_4_NO_3_, 20% PEG 3 350; for STING/MD1203: 100 mM Tris pH 8.0, 20% PEG 8.000, 200 mM LiCl; for STING/MD1202D: 100 mM HEPES pH 7.5, 20% PEG 4.000, 200 mM NaCl). The crystals grew within a range of 24 to 72 hours at room temperature. Upon harvest, the crystals were cryo-protected in the well solution supplemented with 20% glycerol (except for crystals of poxin/MD1203 that did not require cryo-protection) and flash cooled in liquid nitrogen. The datasets were collected from single crystals on the BL14.1 beamline at the BESSY II electron storage ring operated by the Helmholtz-Zentrum Berlin [29].

The data were integrated and scaled using XDSapp v3.1.9 [30, 31]. Structures of the MpxV poxin and human STING proteins in complexes with MD1203 or MD1202D were solved by molecular replacement using the structures of vaccinia virus poxin (pdb entry 6EA6, [10]) and the STING/cGAMP complex (pdb entry 4KSY, [32]) as search models, respectively. Initial models were obtained with Phaser v2.8.3 [33]. The models were further improved using automatic model refinement with the phenix.refine tool [34] from the Phenix package v1.20.1-4487 [35] and manual model building with Coot v0.9.8.7 [36]. Geometrical restraints for the ligands were generated with Grade2 v1.3.1 (Global Phasing Ltd.). Statistics for data collection and processing, structure solution and refinement were calculated with the phenix.table_one tool and are summarized in **Table S1**. Structural figures were generated with the PyMOL Molecular Graphics System v2.5.4 (Schrödinger, LLC). The atomic coordinates and structural factors were deposited in the Protein Data Bank (https://www.rcsb.org).

## Disccussion

In this study, our aim was to prepare cGAMP analogs that would be resistant against the viral endonuclease poxin, which specifically cleaves cGAMP. So far only few analogs of cGAMP, phosphorothioate and dideoxy cGAMP, were described as poxin resistant [10, 12]. The ribose 2’-hydroxyl group is essential for poxin mediated cleavage of cGAMP. It acts as a nucleophile induced by the amino group of poxin’s Lys142 and attacks the bridging phosphate of the 3′-5′ phosphate. This subsequently leads to the cleavage of the bridge and, via 2′-3′ phosphate intermediate, ultimately leads to the linear dinucleotide [10]. However, simple dideoxy cGAMP analogs have certain disadvantages.

compared to the canonical 2′3′cGAMP as well as the difluorinated cGAMP analogue. While the dideoxy analogues perform exceptionally well in assays using digitonin-permeabilized cells, they display significantly lower affinity toward the STING protein in differential scanning fluorimetry (DSF) assays. Furthermore, they show reduced activity on the REF allelic form of STING and display notably decreased performance in standard, unpermeabilized cells [14, 16]. We speculated that by using carba-CDNs with the hydroxyl groups substituted by fluorine atoms, we could create an exceptionally active STING agonist. This agonist would effectively target all STING isoforms, resist poxin-induced cleavage, and exhibit stability against nucleoside hydrolases. We prepared two such compounds: a CDN phosphate, MD1203, and a phosphorothioate, MD1202D. Both of these compounds indeed demonstrated outstanding biological activities against the wild-type STING and its HAQ, AQ, and Q allelic forms. Additionally, MD1202D outperformed the ex-clinical trials candidate ADU-S100 and dideoxy-cGAMP by an order of magnitude against the REF allelic form (**Table 1**). Intriguingly, the conformation of poxin when bound to MDs is almost identical to the conformation of unliganded poxin (Fig. S2), implying that cGAMP can induce conformational changes that the MD compounds cannot.

## Conclusion

These non-hydrolysable, stable cGAMP analogs possess significant potential as antiviral compounds. They effectively bypass viruses’ defense mechanisms involving poxin nucleases that aim to prevent STING pathway activation. The compounds presented here demonstrate remarkable biological activity towards STING. However, further research is needed, including enhancement of cell-membrane permeation, which represents a challenge for all charged molecules. Alternatively, the design of compounds derived from CDNs that no longer bind to STING but display a high affinity for poxins is another approach. This strategy could selectively hinder viral activity while leaving the STING pathway unaffected in uninfected cells.

## Acknowledgment

We thank the Helmholtz-Zentrum Berlin für Materialien und Energie for the allocation of synchrotron radiation beamtime. We are grateful to Dr. Gert Weber and Dr. Frank Lennartz for assistance during crystallographic data collections. This research was funded by the project the National Institute Virology and Bacteriology (Programme EXCELES, Project No. LX22NPO5103) - Funded by the European Union - Next Generation EU. RVO: 61388963 is also acknowledged.

## Author Contribution

M.K., M.D., V.D., A.E., M.S., K.C., and D.C. performed experiments. M.K., K.C., and E.B. analyzed data. M.K., R.N., and E.B. wrote the manuscript. R.N. and E.B. conceived the project and obtained funding.

## Notes

### Competing Interest Statement

The authors have declared no competing interest.

