## Supplementary Information for "Fluorinated cGAMP analogs, which act as STING agonists and are not cleavable by poxins: structural basis of their function"

### Materials and Methods

***Synthesis*** – Unless stated otherwise, all solvents were evaporated at 40 °C at 2 kPa, and the compounds were dried at 30 °C at 2 kPa. Starting compounds and reagents were purchased from commercial suppliers (Sigma-Aldrich, Fluorochem, Acros Organics, Carbosynth) and used without further purification. Acetonitrile was dried using activated 3A molecular sieves. Analytical Thin-Layer Chromatography (TLC) was performed on silica gel-precoated aluminum plates with a fluorescent indicator (Merck 60 F254). Column chromatography (both normal and reverse phase) was performed on a 40-60 µm silica gel using an ISCO flash chromatography system. Purity of the final compounds was determined by UPLC MS and was 95% or higher.

Analytical High-Performance Liquid Chromatography (HPLC), mass spectra, UV absorbance and compound purity were measured on a Waters Ultra-High Performance Liquid Chromatography-Mass Spectrometry (UPLC-MS) system consisting of a Waters UPLC H-Class Core System, a UPLC photodiode array (PDA) detector and a Waters SQD2 or QDa mass spectrometer. The MS method used was electrospray ionization (ESI)+ and/or ESI-, cone voltage = 15 V, mass detector range 200–1000 Da. Two sets of HPLC conditions were used as indicated: (a) C18 (column: Waters Acquity UPLC BEH C18 column, 1.7 mm, 2.1 × 100 mm; LC method: $H_2O$/ACN, 0.1% formic acid as a modifier, gradient 0–100 %, run length 7 min, flow 0.5 ml/min) and (b) HILIC (column: SeQuant ZIC-pHILIC, 5 µm, polymeric, 50 x 2.1 mm; LC method: ACN/0.01M aqueous ammonium acetate gradient 10–60 %, run length 7 min, flow 0.3 ml/min).

The final CDNs were purified by semipreparative HPLC (Luna, 5 µm, C18 250 x 21 mm) using triethylammonium bicarbonate (TEAB) as a modifier. TEAB was removed from the collected fractions by 3 cycles of co-evaporation with methanol.

NMR spectra were recorded on Bruker Avance III HD machines ($^1$H at 400 or 600 MHz) using a solvent signal as a reference. *Tert*-butyl alcohol was used as an internal standard in $D_2O$ solutions. Chemical shifts (δ) and coupling constants (*J*) were expressed in ppm and Hz, respectively. All structures were confirmed, and $^1$H and 13C signals were assigned by combining 1D and 2D NMR (H,H-COSY, H,C-HSQC, H,C-HMBC) techniques and using standard pulse programs from the library of the spectrometer; gradient selection was used in the 2D experiments. High resolution mass spectra were measured on an LTQ Orbitrap XL using electrospray ionization (HRMS ESI).

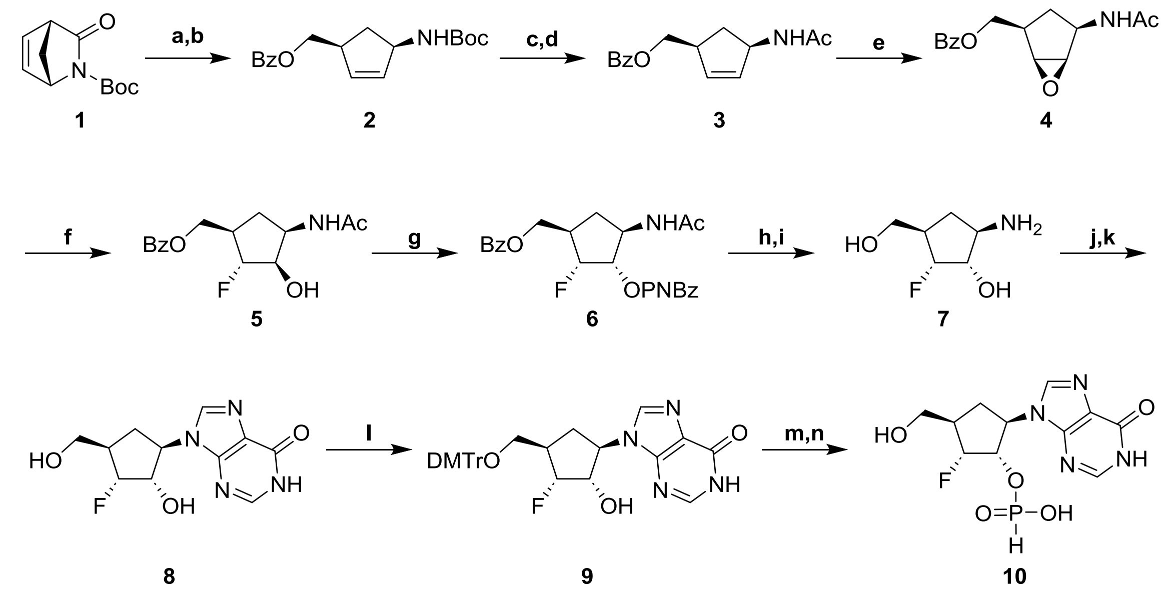

Scheme S1: Synthesis of the key monomer. Reagents and conditions: a) $NaBH_4$, THF-MeOH, 0 °C to r.t., 1 h; b) BzCl, DMAP, pyridine, DCM, 0 °C to r.t., 12 h; c) TFA, DCM, 0 °C to r.t., 16 h; d) $Ac_2O$, pyridine, 0 °C to r.t., 1 h; e) mCPBA, DCM, 0 °C to r.t., 14 h; f) HF-pyridine, DCM, 0 °C to r.t., 10 h; g) 4-nitrobenzoic acid, DIAD, $PPh_3$, toluene, 0 °C to r.t., 24 h; h) $NH_3$-MeOH, r.t. 24 h; i) HCl, EtOH, 80 °C, 24 h; j) *N*-(4,6-dichloropyrimidin-5-yl)formamide, DIPEA, nBuOH, 145 °C, 28 h; k) 80 % formic acid, 55 °C, 16 h; l) DMTrCl, pyridine, 0 °C to r.t., 15 h; m) diphenyl phosphite, pyridine, r.t. 20 min, then TEA, water, r.t. 20 min; n) DCA, TES, DCM, r.t. 30 min.

((1*S*,4*R*)-4-((*tert*-Butoxycarbonyl)amino)cyclopent-2-en-1-yl)methyl benzoate (**2**):

To an ice-cooled solution of *tert*-butyl (1*R*,4*S*)-3-oxo-2-azabicyclo[2.2.1]hept-5-ene-2-carboxylate **1** (8.16 g, 39 mmol) in THF-methanol mixture (210 mL, 9:1) sodium borohydride (2.99 g, 78 mmol) was added in five portions during 30 minutes and the reaction mixture was stirred for 1 h at 0 °C and then further 1 h at room temperature. Reaction mixture was evaporated to 1/3 of its original volume and then partitioned between ethyl acetate (800 mL) and saturated aqueous $NaHCO_3$ (250 mL). Organic phase was then washed with brine (250 mL), dried over sodium sulfate and evaporated. Residue was co-evaporated with toluene (250 mL) and re-dissolved in dichloromethane (290 mL). Pyridine (6.3 mL, 78 mmol) and DMAP (100 mg) were added and the reaction mixture was cooled down to 0 °C. Benzoyl chloride (6 mL, 46.8 mmol) was added dropwise during 10 minutes. Reaction mixture was allowed to warm to room temperature, stirred overnight and evaporated. The residue was then partionated between ethyl acetate (800 mL) and saturated aqueous $NaHCO_3$ (250 mL). Organic phase was then washed with brine (250 mL), dried over sodium sulfate and evaporated. Product was isolated by flash column chromatography (ethyl acetate in cyclohexane 0 → 35%) to afford **2** (11.32 g, 91 %): $^1$H NMR (400 MHz, DMSO-$d_6$) δ 8.05 – 7.90 (m, 2H), 7.73 – 7.62 (m,

1H), 7.58 – 7.50 (m, 2H), 6.99 (d, *J* = 7.9 Hz, 1H), 5.84 (dt, *J* = 5.7, 2.0 Hz, 1H), 5.75 (dt, *J* = 5.8, 2.2 Hz, 1H), 4.54 (d, *J* = 8.0 Hz, 1H), 4.23 (dd, *J* = 6.7, 1.2 Hz, 2H), 3.00 (ddd, *J* = 8.5, 5.4, 2.0 Hz, 1H), 2.42 (dt, *J* = 13.2, 8.2 Hz, 1H), 1.39 (s, 10H); $^{13}$C NMR (101 MHz, DMSO-$d_6$) δ 165.89, 155.13, 134.45, 133.45, 132.85, 129.93, 129.34, 128.89, 77.74, 67.88, 56.02, 43.63, 34.17, 28.43; ESI MS m/z (%): 340.1 (100) [M+Na]; HRMS ESI ($C_{18}H_{23}NO_4Na$) calculated 340.15193; found: 340.15178.

((1*S*,4*R*)-4-Acetamidocyclopent-2-en-1-yl)methyl benzoate (**3**):

To an ice-cooled solution of **2** (11.2 g, 35.3 mmol) in dichloromethane (148 mL), TFA (14.8 mL) was added dropwise (15 minutes). Reaction mixture was allowed to warm to room temperature and was stirred for another 16 hours. Volatiles were evaporated and the residue was co-evaporated with acetonitrile (3 x 150 mL). Oily residue was dissolved in acetonitrile (100 mL) and pyridine (50 mL) and resulting solution was cooled down in an ice bath. Acetic anhydride (6.7 mL, 71 mmol) was added during 5 minutes. Reaction mixture was stirred for another hour and then evaporated, re-dissolved in ethyl acetate (800 mL). The organic phase was washed with saturated aqueous $NaHCO_3$ (250 mL), water (250 mL) and brine (250 mL), dried over sodium sulfate and evaporated. Product was isolated on flash column chromatography (ethyl acetate) to afford **3** (8.58 g, 94 %): $^1$H NMR (400 MHz, DMSO-$d_6$) δ 8.04 – 7.97 (m, 2H), 7.95 (d, *J* = 7.7 Hz, 1H), 7.72 – 7.62 (m, 1H), 7.59 – 7.47 (m, 2H), 5.90 (dt, *J* = 5.6, 2.0 Hz, 1H), 5.75 (dt, *J* = 5.6, 2.2 Hz, 1H), 4.76 (dtt, *J* = 10.1, 6.1, 2.0 Hz, 1H), 4.35 – 4.17 (m, 2H), 3.03 (tddt, *J* = 8.5, 6.5, 4.3, 2.1 Hz, 1H), 2.44 (dt, *J* = 13.3, 8.3 Hz, 1H), 1.78 (s, 3H), 1.34 (dt, *J* = 13.3, 6.3 Hz, 1H); $^{13}$C NMR (101 MHz, DMSO-$d_6$) δ 168.64, 165.90, 134.17, 133.51, 133.34, 129.90, 129.34, 128.94, 67.97, 54.46, 43.72, 34.34, 22.74; ESI MS m/z (%): 282.1 (100) [M+Na]; HRMS ESI ($C_{15}H_{17}NO_3Na$) calculated 282.11007; found: 282.11015.

((1*S*,2*R*,4*R*,5*R*)-4-Acetamido-6-oxabicyclo[3.1.0]hexan-2-yl)methyl benzoate (**4**):

A solution of **3** (8.58 g, 33 mmol) in dichloromethane (410 mL) was cooled to 0 °C, *m*-chloroperoxybenzoic acid (mCPBA, 75%, 11.18 g, 48.6 mmol) was added and the reaction mixture was allowed to warm to room temperature and then stirred for 14 h. Reaction mixture was evaporated to about 1/3 of its original volume and diluted with ethyl acetate (1.2 L). This solution was washed with saturated aqueous sodium carbonate (5 x 500 mL), dried over sodium sulfate, and evaporated. The obtained solid was mixed with diethyl ether and resulting suspension was sonicated for 20 minutes. The solid was collected and dried *in vacuo* to obtain **4** (8.1 g, 89 %): $^1$H NMR (400 MHz, DMSO-$d_6$) δ 8.10 (d, *J* = 7.5 Hz, 1H), 8.05 – 7.98 (m, 2H), 7.72 – 7.64 (m, 1H), 7.59 – 7.49 (m, 2H), 4.36 (dd, *J* = 10.8, 6.3 Hz, 1H), 4.32 – 4.16 (m, 2H), 3.56 (dd, *J* = 2.9, 1.2 Hz, 1H), 3.50 (dd, *J* = 2.9, 1.3 Hz, 1H), 2.46 (ddt, *J* = 7.8, 2.6, 1.3 Hz, 1H), 1.93 – 1.84 (m, 1H), 1.82 (s, 3H), 0.92 (dt, *J* = 12.4, 9.8 Hz, 1H); $^{13}$C NMR (101 MHz, DMSO-$d_6$) δ 169.45,

165.81, 133.54, 129.81, 129.40, 128.92, 64.71, 57.61, 55.68, 49.91, 37.70, 27.00, 22.59; ESI MS m/z (%): 298.1 (100) [M+Na]; HRMS ESI ($C_{15}H_{17}NO_4Na$) calculated 298.10498; found: 298.10492.

((1*R*,2*R*,3*R*,4*R*)-4-Acetamido-2-fluoro-3-hydroxycyclopentyl)methyl benzoate (**5**):

Compound **4** (4 g, 14.53 mmol) was co-evaporated with toluene (60 mL) and dichloromethane (60 mL), dissolved in dichloromethane (80 mL) in a polypropylene flask under argon atmosphere. Reaction mixture was cooled down to 0 °C, HF/pyridine (30% HF, 8.4 mL) was added dropwise and the reaction mixture was left to slowly warm to room temperature and then stirred for 10 h. Reaction mixture was poured into a mixture of ice / saturated $NaHCO_3$ (800 mL) and water phase was extracted with dichloromethane (4 x 400 mL). Combined organic phases were dried with sodium sulfate and evaporated. Product was isolated by column chromatography ($CHCl_3$:acetone 3:2) to afford **5** (3.69 g, 86 %): $^1$H NMR (400 MHz, DMSO-$d_6$) δ 8.12 – 7.95 (m, 2H), 7.74 (d, *J* = 7.8 Hz, 1H), 7.73 – 7.62 (m, 1H), 7.60 – 7.48 (m, 2H), 5.41 (d, *J* = 4.2 Hz, 1H), 4.73 (ddd, *J* = 51.1, 3.5, 2.2 Hz, 1H), 4.43 – 4.25 (m, 2H), 4.23 – 4.09 (m, 1H), 4.05 – 3.93 (m, 1H), 2.49 – 2.29 (m, 1H), 2.20 – 2.04 (m, 1H), 1.82 (s, 3H), 1.48 (ddd, *J* = 12.6, 10.6, 9.2 Hz, 1H); $^{13}$C NMR (101 MHz, DMSO-$d_6$) δ 169.28, 165.83, 133.58, 129.77, 129.35, 128.94, 99.60 (d, *J* = 179.6 Hz), 73.71 (d, *J* = 25.3 Hz), 65.64 (d, *J* = 6.1 Hz), 50.54, 41.25 (d, *J* = 22.1 Hz), 30.21 (d, *J* = 2.8 Hz), 22.78; $^{19}$F NMR (376 MHz, DMSO-$d_6$) δ -177.09 (dddd, *J* = 51.0, 30.4, 12.4, 2.0 Hz); ESI MS m/z (%): 318.1 (100) [M+Na]; HRMS ESI ($C_{15}H_{18}NO_4FNa$) calculated 318.11121; found: 318.11126.

(1*S*,2*R*,3*R*,5*R*)-5-Acetamido-3-((benzoyloxy)methyl)-2-fluorocyclopentyl 4-nitrobenzoate (**6**):

Compound **5** (3.68 g, 12.46 mmol) was co-evaporated with toluene (2 x 100 mL), dissolved in toluene (185 mL) and then *p*-nitrobenzoic acid (6.26 g, 37.4 mmol) and triphenylphosphine (9.8 g, 37.4 mmol) were added. Reaction mixture was cooled down to 0 °C and DIAD (7.36 mL, 37.4 mmol) was added dropwise. Reaction mixture was left to slowly warm to room temperature and then stirred for 24 h, diluted with ethyl acetate (500 mL) and washed with saturated $NaHCO_3$ (4 x 250 mL). Organic phase was dried with sodium sulfate and evaporated. Product was isolated by column chromatography (toluene:acetone 4:1) to afford **6** (2.1 g, 41 %): $^1$H NMR (400 MHz, DMSO-$d_6$) δ 8.37 – 8.27 (m, 2H), 8.27 – 8.18 (m, 2H), 8.12 (d, *J* = 7.8 Hz, 1H), 7.94 (dd, *J* = 8.3, 1.4 Hz, 2H), 7.65 – 7.60 (m, 1H), 7.52 – 7.35 (m, 2H), 5.36 (ddd, *J* = 13.9, 5.9, 3.0 Hz, 1H), 5.14 (ddd, *J* = 50.9, 5.3, 3.0 Hz, 1H), 4.60 – 4.33 (m, 3H), 2.75 – 2.55 (m, 1H), 2.40 – 2.23 (m, 1H), 1.74 (s, 3H), 1.76 – 1.67 (m, 1H); $^{13}$C NMR (101 MHz, DMSO-$d_6$) δ 169.35, 165.79, 163.54, 150.53, 134.84, 133.53, 131.05, 129.63, 129.29, 128.84, 123.86, 97.38 (d, *J* = 182.6 Hz), 77.34 (d, *J* = 27.4 Hz), 64.81 (d, *J* = 3.0 Hz), 48.34, 40.78 (d, *J* = 21.2 Hz), 30.10, 22.48; $^{19}$F NMR (376 MHz, DMSO-$d_6$) δ -180.74 (ddd, *J* = 51.0, 28.3, 13.8 Hz); ESI MS m/z (%): 467.2 (100) [M+Na]; HRMS ESI ($C_{22}H_{21}N_2O_7FNa$) calculated 467.12250; found: 467.12223.

9-((1*R*,2*S*,3*R*,4*R*)-3-Fluoro-2-hydroxy-4-(hydroxymethyl)cyclopentyl)-1,9-dihydro-6*H*-purin-6-one (**8**):

Compound **6** (1 g, 2.25 mmol) was dissolved in methanolic ammonia (10 M, 23 mL) and reaction mixture was stirred 24 h at room temperature and then evaporated. Residue was co-evaporated with ethanol (2 x 50 mL), dissolved in in a mixture of 2M HCl (12.5 ml) and ethanol (12.5 mL) the resulting solution was heated to 95 °C for 24 hours. Reaction mixture was neutralized with solid $NaHCO_3$ and then the solution was applied on a Dowex 50 column ($H^+$ cycle, 200 mL). Column was washed with water and methanol, and crude **7** was then eluted with aqueous ammonia-methanol (1:4). Fractions containing **7** were pooled, evaporated and residue was co-evaporated with ethanol.

Crude **7** (320 mg) was dissolved in *n*-butanol (11 mL) and DIPEA (0.83 ml, 4.8 mmol). To this solution, *N*-(4,6-dichloropyrimidin-5-yl)formamide (555 mg, 2.89 mmol) was added and the reaction mixture was heated in a pressure vessel to 145 °C for 28 hours. Volatiles were evaporated and the residue was purified on a silica gel column (ethyl acetate → ethyl acetate:toluene:acetone:ethanol 17:4:3:1, 0-100%). The obtained solid was dissolved in aqueous HCOOH (80%, 8.5 mL) and heated to 55 °C for 16 hours, evaporated and co-evaporated with ethanol (2 x 25 mL). The residue was then dissolved in ethanol (15 mL) and aqueous ammonia (25%, 5 mL) was added. After 30 minutes the mixture was evaporated, co-evaporated with ethanol (2 x 25 mL) and product was isolated by a reverse-phase FCC ($H_2O$:acetonitrile 0 → 30%) to afford **8** (157 mg, 26 %): $^1$H NMR (400 MHz, DMSO-$d_6$) δ 12.26 (br s, 1H), 8.08 (s, 1H), 8.04 (s, 1H), 5.58 (br s, 1H), 4.98 – 4.85 (m, 2H), 4.78 (ddd, $J$ = 51.2, 3.3, 2.1 Hz, 1H), 4.26 – 4.09 (m, 1H), 3.53 (qd, $J$ = 10.6, 6.1 Hz, 2H), 2.40 – 2.17 (m, 2H), 2.06 (td, $J$ = 11.9, 9.1 Hz, 1H); $^{13}$C NMR (101 MHz, DMSO-$d_6$) δ 156.87, 148.70, 145.49, 139.84, 123.85, 99.19 (d, $J$ = 179.4 Hz), 73.78 (d, $J$ = 26.2 Hz), 62.21 (d, $J$ = 5.5 Hz), 55.02 (d, $J$ = 1.6 Hz), 44.64 (d, $J$ = 20.2 Hz), 29.34 (d, $J$ = 3.2 Hz). $^{19}$F NMR (376 MHz, DMSO-$d_6$) δ -176.02 (ddd, $J$ = 51.2, 30.9, 12.2 Hz); ESI MS m/z (%): 291.1 (100) [M+Na]; HRMS ESI ($C_{11}H_{13}N_4O_3FNa$) calculated 291.08639; found: 291.08645.

9-((1*R*,2*S*,3*R*,4*R*)-4-((Bis(4-methoxyphenyl)(phenyl)methoxy)methyl)-3-fluoro-2-hydroxycyclopentyl)-1,9-dihydro-6*H*-purin-6-one (**9**):

Compound **8** (148 mg, 0.55 mmol) was azeotroped with pyridine (3 x 15 mL), dissolved in pyridine (10 mL) and DMTrCl (206 mg, 0.61 mmol) was added in one portion at 0 °C. Reaction mixture was allowed to warm to room temperature and stirred for 15 h, then quenched with saturated aqueous $NaHCO_3$ (3 mL) and evaporated. Residue was taken up in ethyl acetate (200 mL) and washed with saturated aqueous solution of $NaHCO_3$ (100 mL). Organic phase was dried over sodium sulfate and evaporated. Product was purified on FCC (ethyl acetate → ethyl acetate:acetone:ethanol:water 20:3:1.2:0.8, 0-100%) to afford **9** (284 mg, 90 %): $^{1}$H NMR (400 MHz, DMSO-$d_6$) δ 12.27 (s, 1H), 8.03 (s, 1H), 8.02 (s, 1H), 7.47 – 7.38 (m, 2H), 7.37 – 7.14 (m, 7H), 6.98 – 6.84 (m, 4H), 5.56 (d, *J* = 4.8 Hz, 1H), 5.04 – 4.67 (m, 2H), 4.30 – 4.11 (m, 1H), 3.74 (s, 6H), 3.17 (d, *J* = 6.9 Hz, 2H), 2.36 (dt, *J* = 12.6, 7.5 Hz, 2H), 2.14 – 2.00 (m, 1H); $^{13}$C NMR (101 MHz, DMSO) δ 158.23, 156.84, 148.65, 145.41, 145.16, 140.05, 135.85, 129.84, 128.04, 127.83, 126.84, 123.92, 113.39, 99.84 (d, *J* = 181.1 Hz), 85.54, 73.63 (d, *J* = 25.8 Hz), 64.17 (d, *J* = 4.0 Hz), 55.21, 54.81 (d, *J* = 1.8 Hz), 45.85, 42.40 (d, *J* = 21.3 Hz), 29.74 (d, *J* = 2.6 Hz); $^{19}$F NMR (376 MHz, DMSO-$d_6$) δ -175.64 (ddd, *J* = 51.5, 30.1, 12.7 Hz); ESI MS m/z (%): 593.3 (100) [M+Na]; HRMS ESI ($C_{32}H_{31}N_4O_5FNa$) calculated 593.21707; found: 593.21667.

(1*S*,2*R*,3*R*,5*R*)-2-Fluoro-3-(hydroxymethyl)-5-(6-oxo-1,6-dihydro-9*H*-purin-9-yl)cyclopentyl hydrogen phosphonate, triethylammonium salt (**10**):

Compound **9** (262 mg, 0.46 mmol) was azeotroped with pyridine (2 x 15 mL), dissolved in pyridine (10 mL) and diphenyl phosphite (85%, 312 µL, 1.39 mmol) was added in one portion. After stirring at ambient temperature for 20 minutes, TEA (0.69 mL) was added followed by water (0.69 mL) and the reaction was stirred for further 20 minutes. Resulting solution was diluted with dichloromethane (200 mL) and washed with saturated aqueous solution of $NaHCO_3$ (75 mL). Water phase was extracted with dichloromethane (2 x 150 mL). Combined organic phases were dried over sodium sulfate and evaporated. Resulting intermediate was purified on FCC (MeOH in DCM (modifier 1 % $Et_3N$) 0 to 50%). To a solution of the obtained intermediate in dichloromethane (10 mL) was added water (200 µL, 11.1 mmol) followed by a solution of DCA (342 µL, 4.15 mmol) in dichloromethane (4 mL). Reaction mixture was stirred at ambient temperature for 30 minutes, after which it was quenched with triethylsilane (3.6 mL). Reaction mixture was then stirred for 1 h and then pyridine (4 ml) was added and all volatiles were evaporated. Purification on reverse phase FCC (ACN in water, 0-30%) afforded **10** (144 mg, 72 %): $^{1}$H NMR (400 MHz, DMSO-$d_6$) δ 8.08 (s, 1H), 8.02 (s, 1H), 6.16 (d, *J* = 582.5 Hz, 1H), 5.31 (s, 1H), 5.07 – 4.88 (m, 2H), 4.60 (td, *J* = 10.4, 4.8 Hz, 1H), 3.52 (t, *J* = 5.9 Hz, 2H), 3.43 – 3.30 (m, 4H), 2.44 – 2.18 (m, 2H), 2.08 – 1.98 (m, 1H), 1.23 (t, *J* = 7.3 Hz, 6H); $^{13}$C NMR (101 MHz, DMSO) δ 156.83, 148.60, 145.49, 139.64, 123.80, 98.16 (d, *J* = 178.8 Hz), 75.45 (dd, *J* = 28.0, 4.7 Hz), 62.73, 62.40 (d, *J* = 6.4 Hz), 54.58 (d, *J* = 4.9 Hz), 52.18, 44.97 (d, *J* = 20.3 Hz), 29.84, 7.35; $^{31}$P NMR (162 MHz, DMSO-$d_6$) δ 2.15; $^{19}$F NMR (376 MHz, DMSO-$d_6$) δ -174.05 (ddd, *J* = 50.4, 32.0, 10.6 Hz); ESI MS m/z (%): 355.1 (100) [M+Na]; HRMS ESI ($C_{11}H_{14}N_4O_5PNa$) calculated 355.05781; found: 355.05761.

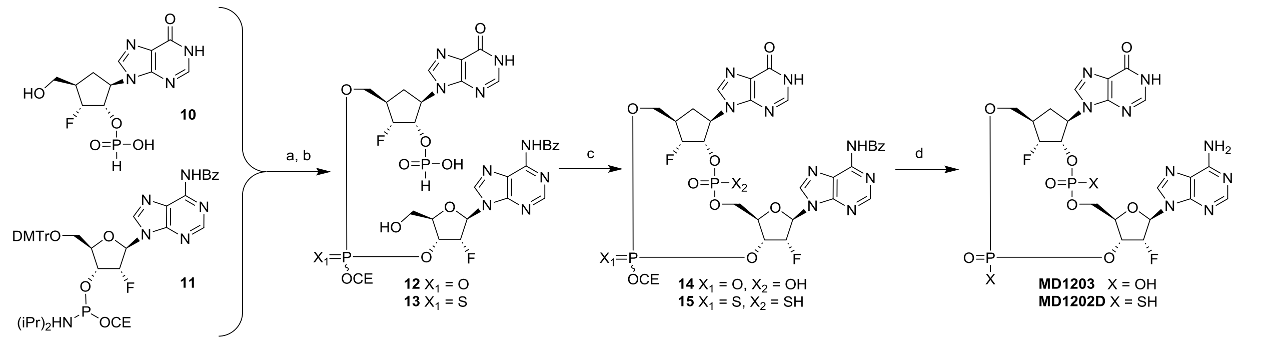


Scheme S2. Synthesis of cyclic dinucleotides. Reagents and conditions: a) py-TFA, *t*BHP, ACN, r.t. 30 min ($X_1$ = O) or py-TFA, 3-((*N*,*N*-dimethylaminomethylidene)amino)-3*H*-1,2,4-dithiazole-5-thione, ACN, r.t. 30 min ($X_1$ = S); b) DCA, TES, DCM, r.t. 30 min; c) DMOCP, $I_2$, pyridine, r.t. 1 h ($X_2$ = OH) or DMOCP, 3*H*-1,2-benzodithiol-3-one, pyridine, r.t. 10 min ($X_2$ = SH); d) $CH_3NH_2$, EtOH, r.t. 3 h.

(1*S*,6*R*,8*R*,9*R*,10*R*,15*R*,17*R*,18*R*)-8-(6-amino-9*H*-purin-9-yl)-9,18-difluoro-3,12-dihydroxy-17-(6-oxo-6,9-dihydro-1*H*-purin-9-yl)-2,4,7,11,13-pentaoxa-3$\lambda^5$ ,12$\lambda^5$ -diphosphatricyclo[13.2.1.0$^{6\ ,10}$ ]octadecane-3,12-dione (**16**)

A mixture of **10** (43 mg, 0.1 mmol) and pyridinium trifluoroacetate (py-TFA, 29 mg, 0.15 mmol) was azeotroped with dry ACN (3 x 3 mL), suspended in dry ACN (1 mL) and stirred overnight in a sealed vessel over activated molecular sieves. In a separate flask, **11** (109 mg, 0.125 mmol, Sigma-Aldrich) was azeotroped with dry ACN (3x 3 mL), dissolved in dry ACN (1 mL) and stirred overnight in a sealed vessel over activated molecular sieves. A solution of phosphoramidite was transferred *via syringe* to the flask with the suspension of **10** with py-TFA and the resulting solution was stirred for 1 hour at ambient temperature. tert-Butyl hydroperoxide (*t*BHP, 5.5M solution in decane, 55 μL, 0.3 mmol) was added and the reaction mixture was stirred for further 30 minutes. Reaction mixture was quenched with $NaHSO_3$ (39% soln. in water, 54 μL, 0.27 mmol), filtered and evaporated. To a solution of the crude tritylated linear dimer in DCM (3 mL) was added water (18 μL, 1 mmol) and a solution of DCA (74 μL, 0.9 mmol) in DCM (3 mL) dropwise. After stirring the reaction mixture for 30 minutes, TES (1.5 mL) was added and the reaction mixture was stirred for further 90 minutes, after which it was quenched by the addition of pyridine (1.5 mL). Volatiles were evaporated and crude **12** (mixture of diastereomers) was codistilled with dry pyridine (3x 3 mL) and used in the next reaction without further purification.

To a solution of crude **12** in pyridine (2 mL) was added DMOCP (65 mg, 0.35 mmol) and reaction mixture was stirred at ambient temperature for 1 hour. Water (59 μL, 0.35 mmol) was added followed by iodine (34 mg, 0.13 mmol) and reaction mixture was stirred for 10 minutes, after which it was cooled down to 0 °C and quenched by the addition of $NaHSO_3$ (39% soln. in water, 49 □L, 0.25 mmol). Purification on reverse phase FCC (ACN in 50 mM aqueous $NH_4HCO_3$ 0-70%) afforded **14** (mixture of diastereomers).

A solution of **14** (20 mg) in $CH_3NH_2$ (33% in ethanol, 1 mL) was stirred at ambient temperature for 3 hours. Volatiles were evaporated and the residue was purified on preparative HPLC (ACN in 0.1M TEAB, 0-30%). Appropriate fractions were pooled, evaporated, codistilled with water (3x 20 mL) and methanol (3x 20 mL), dissolved in water (10 mL) and slowly passed through a 10 mL column of Dowex 50 (Na+ cycle). Freeze-drying the eluent afforded sodium salt of Compound **16** (16 mg, 24 %). HPLC retention time (C18, min): 2.23 min; $^{1}$H NMR (600 MHz, $D_2O$) δ 8.30 (s, 1H), 8.24 (s, 1H), 8.23 (s, 1H), 8.18 (s, 1H), 6.44 (dd, *J* = 17.6, 1.5 Hz, 1H), 5.57 (ddd, *J* = 51.6, 4.6, 1.5 Hz, 1H), 5.21 (dd, *J* = 48.5, 4.0 Hz, 1H), 5.17 (m, 1H), 5.02 (dddd, *J* = 20.2, 8.5, 6.8, 4.6 Hz, 1H), 4.85 (td, *J* = 8.4, 4.0 Hz, 1H), 4.50 (dt, *J* = 8.5, 3.3 Hz, 1H), 4.35 (dt, *J* = 12.0, 3.5 Hz, 1H), 4.27 (dt, *J*= 10.4, 3.2 Hz, 1H), 4.18 (ddd, *J* = 12.0, 6.6, 3.4 Hz, 1H), 4.03 (dt, *J* = 10.4, 5.5 Hz, 1H), 2.71 (m, 1H), 2.51 (m, 2H); $^{31}$P NMR (202.4 MHz, $D_2O$) δ 0.40, 0.29; $^{19}$F NMR (376 MHz, $D_2O$) δ -170.55, -197.71.

(1S,6R,8R,9R,10R,15R,17R,18R)-8-(6-amino-9H-purin-9-yl)-9,18-difluoro-17-(6-oxo-6,9-dihydro-1H-purin-9-yl)-3,12-disulfanyl-2,4,7,11,13-pentaoxa-3$\lambda^5$ ,12$\lambda^5$ -diphosphatricyclo[13.2.1.$0^{6,10}$ ]octadecane-3,12-dione (**17**)

A mixture of **10** (43 mg, 0.1 mmol) and pyridinium trifluoroacetate (29 mg, 0.15 mmol) was codistilled with dry MeCN (3x 3 mL), suspended in dry MeCN (1 mL) and stirred overnight in a sealed vessel over activated molecular sieves. In a separate flask, **11** (109 mg, 0.125 mmol, Sigma-Aldrich) was codistilled with dry MeCN (3x 3 mL), dissolved in dry MeCN (1 mL) and stirred overnight in a sealed vessel over activated molecular sieves. A solution of phosphoramidite was transferred *via syringe* to the flask with the suspension of **10** with pyridinium trifluoroacetate (py-TFA) and the resulting solution was stirred for 1 hour at ambient temperature. 3-((*N*,*N*-Dimethylaminomethylidene)amino)-3*H*-1,2,4-dithiazole-5-thione (23 mg, 0.11 mmol) was added and the reaction mixture was stirred for further 30 minutes. Volatiles were evaporated to afford crude tritylated linear dimer, which was dissolved in DCM (3 mL), water was added (18 µL, 1 mmol) and a solution of DCA (74 µL, 0.9 mmol) in DCM (3 mL) was added dropwise. After stirring the reaction mixture for 30 minutes, TES (1.5 mL) was added and the reaction mixture was stirred for further 90 minutes, after which it was quenched by the addition of pyridine (1.5 mL). Volatiles were evaporated and crude **13** was codistilled with dry pyridine (3x 3 mL) and used in the next reaction without further purification.

To a solution of crude **13** in pyridine (2 mL) was added DMOCP (65 mg, 0.35 mmol) and reaction mixture was stirred at ambient temperature for 1 hour. Water (18 µL, 1 mmol) was added followed by 3*H*-

1,2-benzodithiol-3-one (25 mg, 0.15 mmol) and reaction mixture was stirred for 10 minutes. Volatiles were evaporated and product was isolated on reverse phase FCC (MeCN in 50 mM aqueous $NH_4HCO_3$ 0-70%) to afford **15** (mixture of diastereomers).

A solution of **15** (21 mg) in $CH_3NH_2$ (33% in ethanol, 1 mL) was stirred at ambient temperature for 3 hours. Volatiles were evaporatedand the product was purified on preparative HPLC (MeCN in 0.1M TEAB, 0-30%). After repeated freeze-drying of the pooled appropriate fractions to remove TEAB, the desired compound was isolated as four separable diastereomers. **17D** (9 mg, 13 %) was the most abundant and last eluting. HPLC retention time (C18, min): 2.63 min; $^{1}$H NMR (600 MHz, DMSO-$d_6$) δ 8.34 (s, 1H), 8.25 (s, 1H), 8.22 (s, 1H), 8.19 (s, 1H), 6.46 (dd, $J$ = 16.7, 2.0 Hz, 1H), 5.60 (ddd, $J$ = 51.2, 4.6, 2.0 Hz, 1H), 5.25 (dd, $J$ = 48.7, 2.1 Hz, 1H), 5.18-5.16 (m, 2H), 5.08 (dddd, $J$ = 12.1, 9.5, 4.0, 2.1 Hz, 1H), 4.53 (td, $J$ = 7.8, 2.9 Hz, 1H), 4.32 (ddd, $J$ = 10.5, 4.6, 3.2 Hz, 1H), 4.27 (dd, $J$ = 7.3, 2.9 Hz, 2H), 4.08 (ddd, $J$ = 10.5, 7.3, 4.5 Hz, 1H), 2.72 (d, $J$ = 38.0 Hz, 1H), 2.54 (m, 2H); $^{31}$P NMR (202.4 MHz, DMSO-$d_6$) δ 56.60, 56.12; $^{19}$F NMR (470.4 MHz, DMSO-$d_6$) δ -170.51, -197.68.

Copies of $^{1}H$, $^{19}F$ and $^{31}P$ NMP NMR spectra

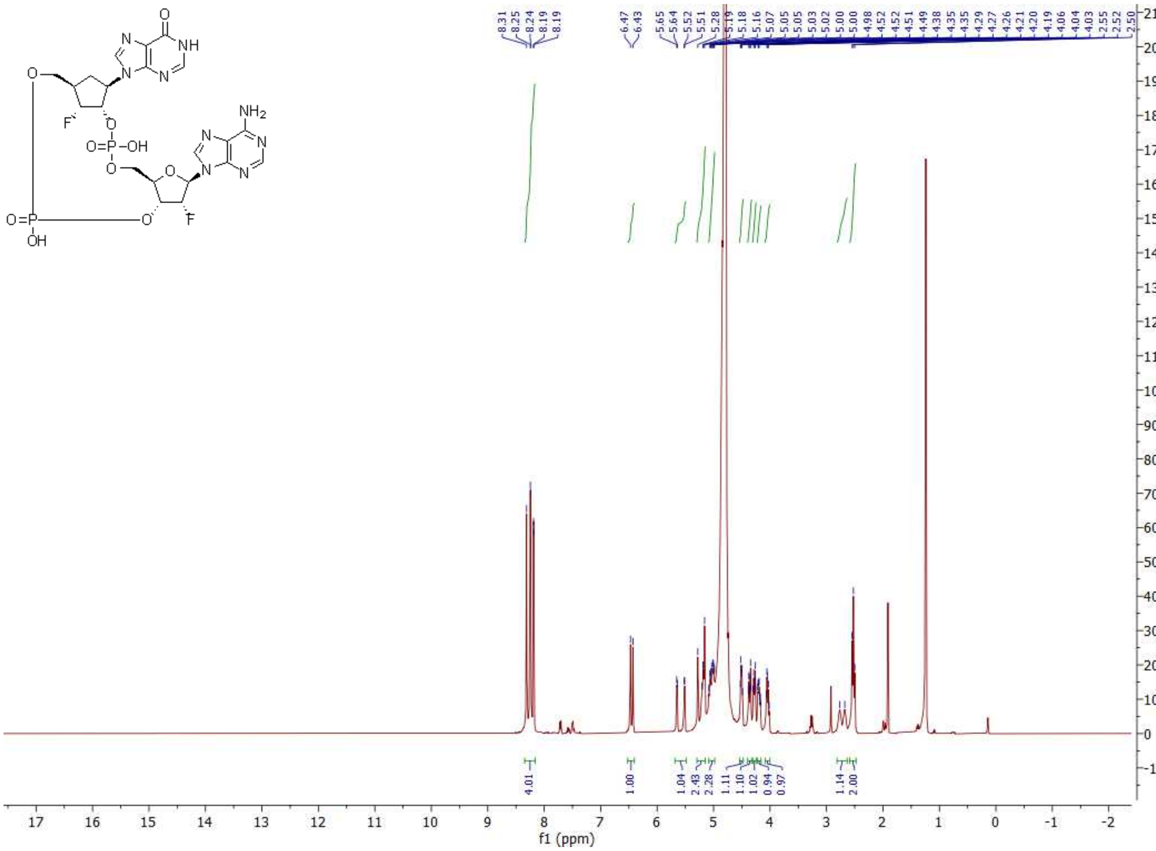

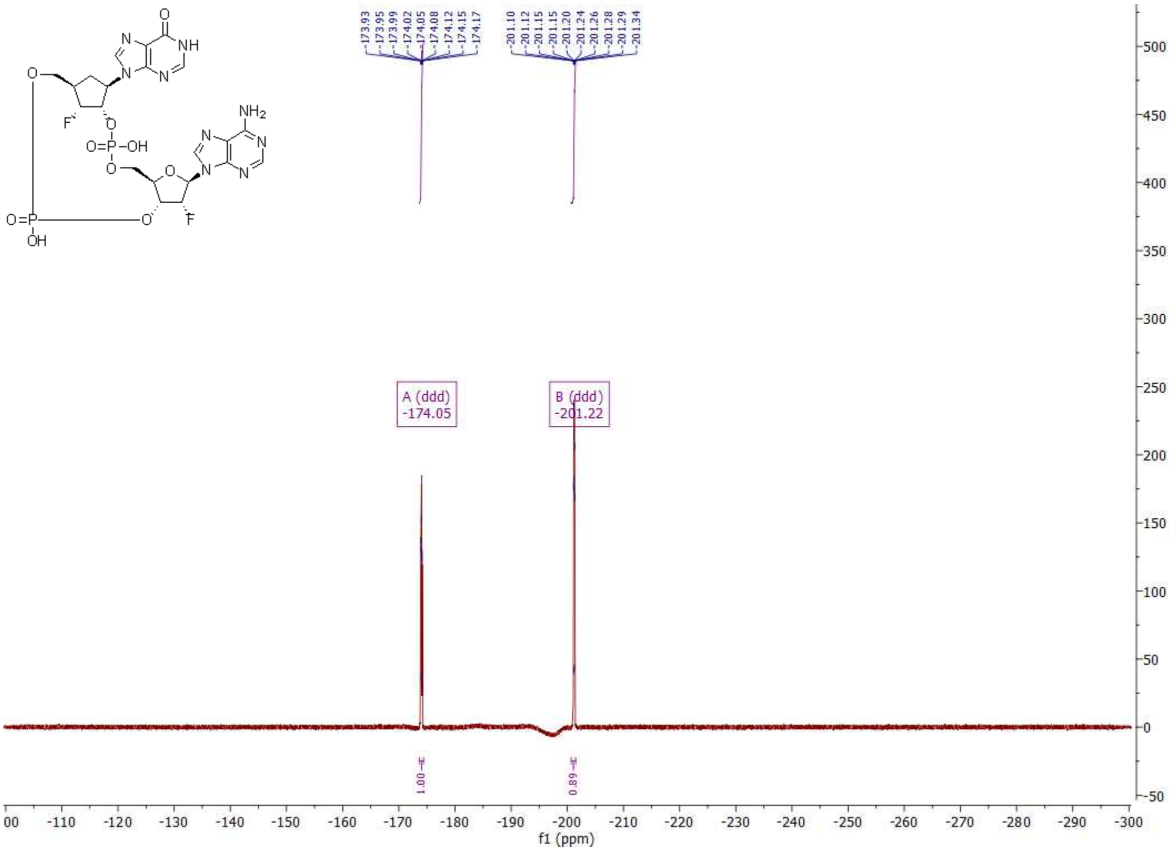
A (ddd)
-174.05
B (ddd)
-201.22
1.00
0.89
f1 (ppm)

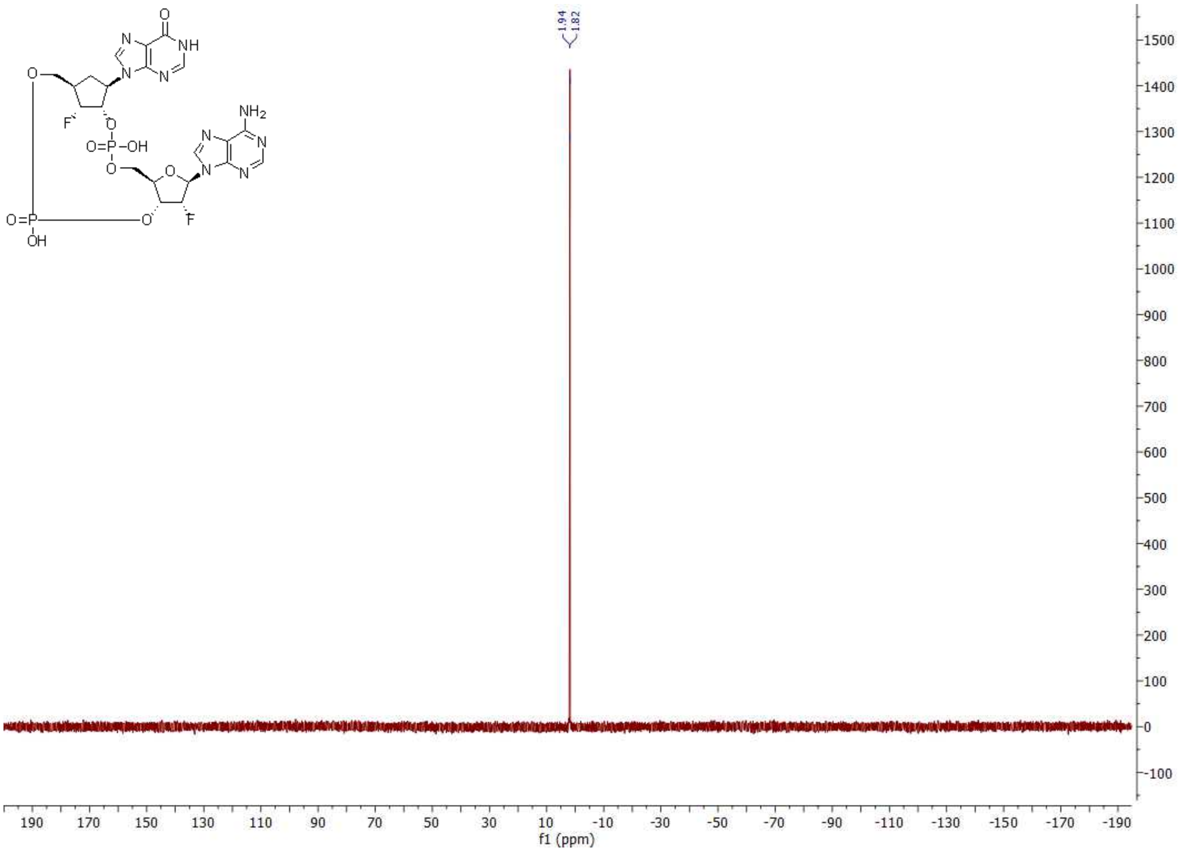
1.94
1.82
f1 (ppm)

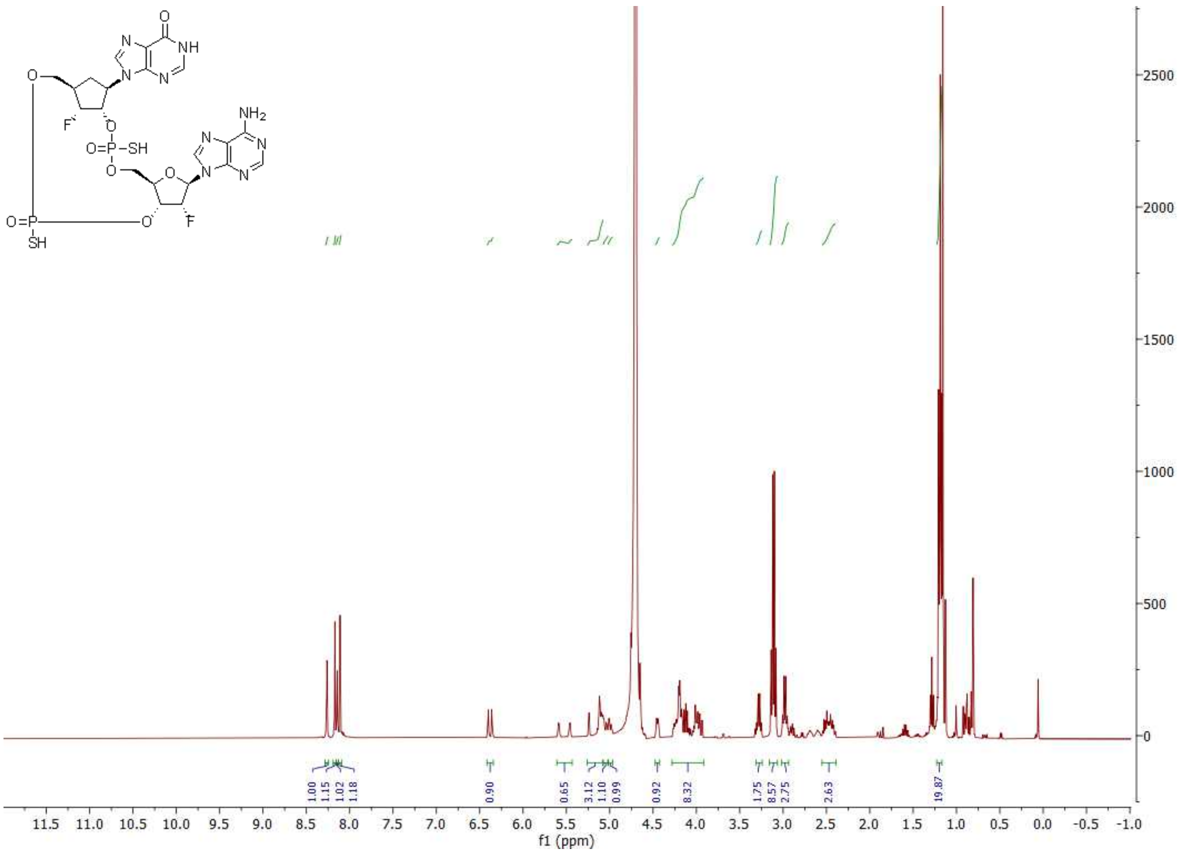
O
NH
N
N
N
NH2
N
N
N
N
O
F
O
O=P-SH
O
O
O=P
SH
O
F
1.00
1.15
1.02
1.18
0.90
0.65
3.12
1.10
0.99
0.92
8.32
1.75
8.57
2.75
2.63
19.87
2500
2000
1500
1000
500
0
11.5 11.0 10.5 10.0 9.5 9.0 8.5 8.0 7.5 7.0 6.5 6.0 5.5 5.0 4.5 4.0 3.5 3.0 2.5 2.0 1.5 1.0 0.5 0.0 -0.5 -1.0
f1 (ppm)

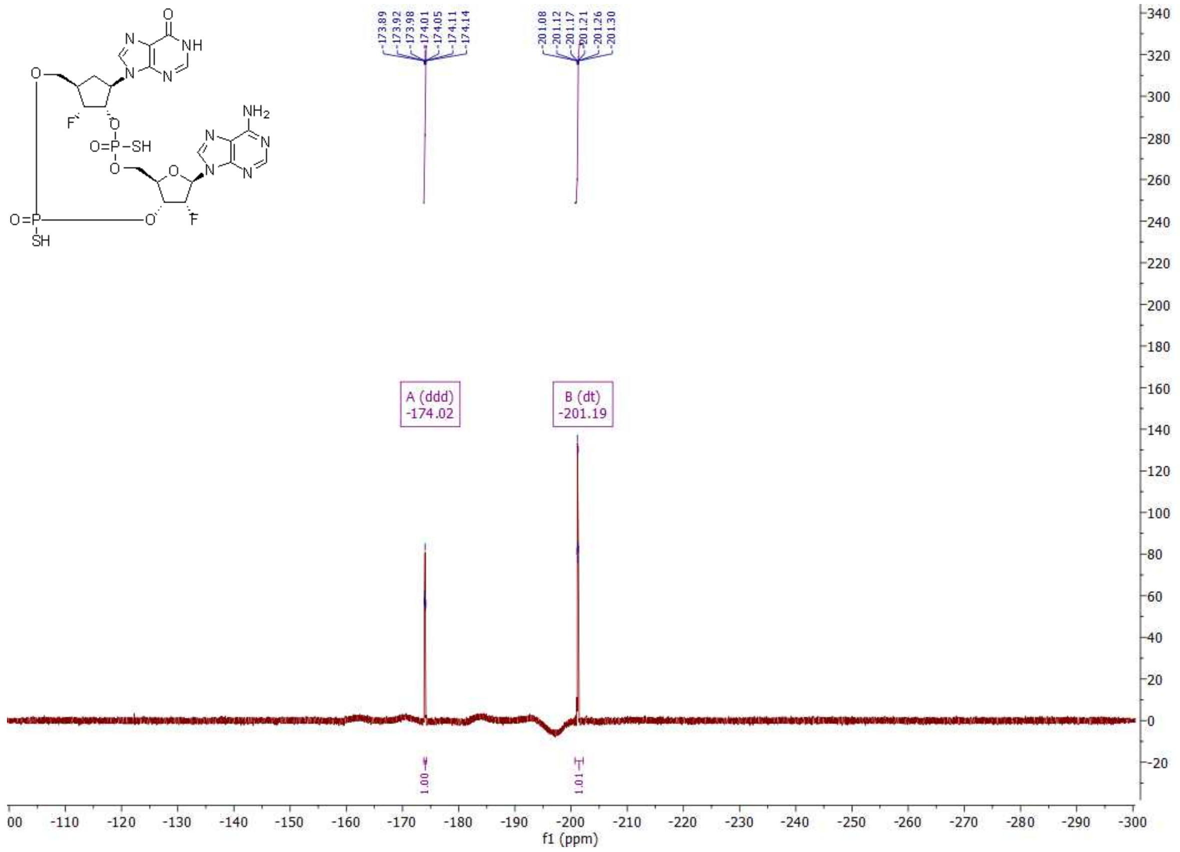
A (ddd)
-174.02
B (dt)
-201.19
1.00
1.01
f1 (ppm)

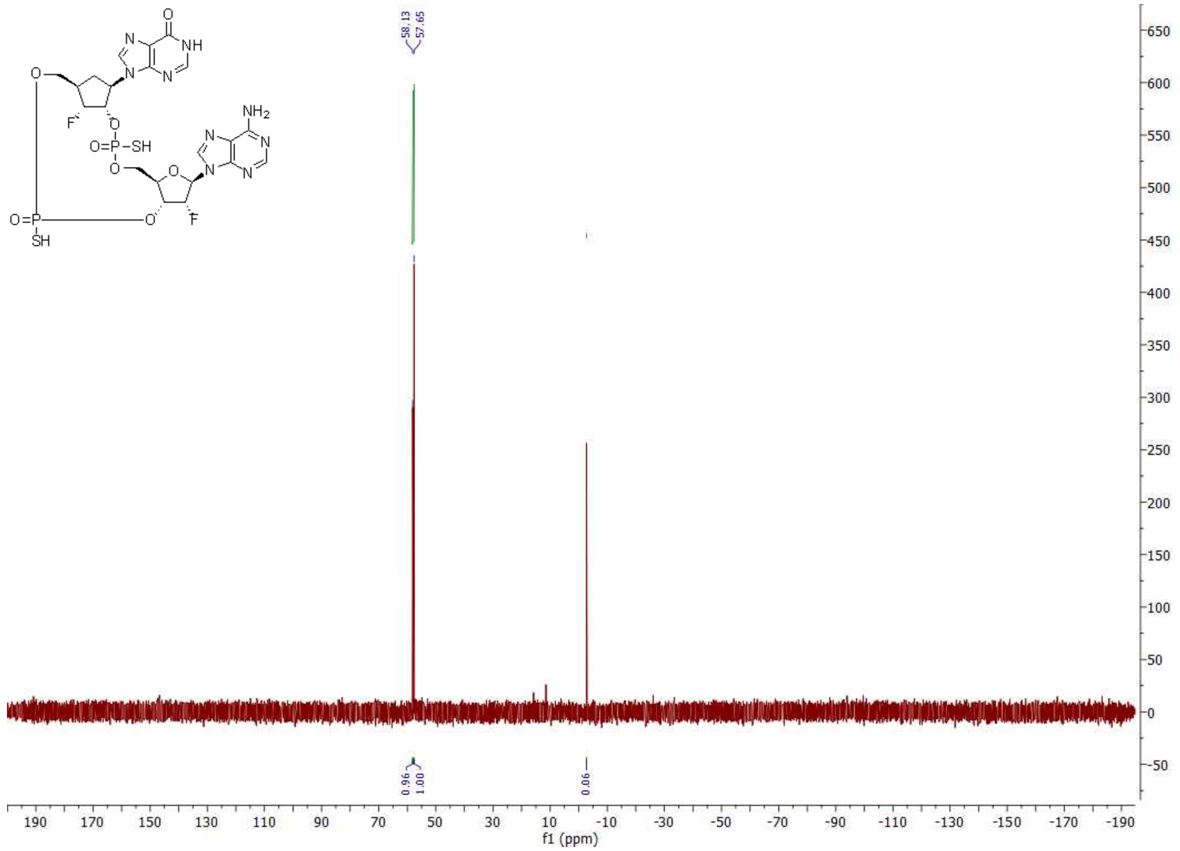
58.13
57.65
0.96
1.00
0.06
f1 (ppm)

| Crystal | | poxin/MD1203 | poxin/MD1202D | STING/MD1203 | STING/MD1202D |
|---|---|---|---|---|---|
| PDB accession code | | 8ORV | 8P44 | 8ORW | 8P45 |
| **Data collection and processing** | | | | | |
| Diffraction source | | BESSY 14.1 | BESSY 14.1 | BESSY 14.1 | BESSY 14.1 |
| Detector | | Dectris Pilatus 6M | Dectris Pilatus 6M | Dectris Pilatus 6M | Dectris Pilatus 6M |
| Wavelength (Å) | | 0.9184 | 0.9184 | 0.9184 | 0.9184 |
| Space group | | C 2 2 $2_1$ | P 1 $2_1$ 1 | C 1 2 1 | C 1 2 1 |
| Cell dimensions | a, b, c (Å) | 54.1 95.5 93.5 | 54.2 91.2 93.1 | 89.4 78.1 35.9 | 89.5 78.0 35.4 |
| | α, β, γ (°) | 90.0 90.0 90.0 | 90.0 90.0 90.0 | 90.0 97.4 90.0 | 90.0 97.6 90.0 |
| Resolution range (Å) | | 33.40 – 1.65 (1.71 – 1.65) | 46.86 - 1.93 (2.00 - 1.93) | 31.68 - 2.95 (3.01 - 2.95) | 35.11 - 3.23 (3.35 - 3.23) |
| No. of total reflections | | 378,583 (33,821) | 463,983 (46,880) | 32,099 (3,178) | 47,406 (2,548) |
| No. of unique reflections | | 29,391 (2,841) | 66,736 (6,613) | 4,988 (494) | 3,917 (370) |
| Completeness (%) | | 99.47 (97.76) | 97.86 (97.66) | 95.11 (93.23) | 99.90 (100.00) |
| Multiplicity | | 12.9 (11.9) | 7.0 (7.1) | 6.4 (6.4) | 12.1 (6.9) |
| Mean I/σ(I) | | 14.98 (0.92) | 8.17 (1.00) | 6.43 (0.77) | 8.20 (1.22) |
| Wilson B factor (Å$^2$) | | 24.19 | 26.56 | 75.27 | 90.47 |
| R-merge | | 0.1307 (2.704) | 0.2053 (2.153) | 0.3852 (2.025) | 0.3004 (1.940) |
| R-meas | | 0.1361 (2.824) | 0.2217 (2.321) | 0.4183 (2.197) | 0.3135 (2.096) |
| CC1/2 (%) | | 99.9 (50.8) | 99.7 (50.2) | 98.5 (45.5) | 99.5 (49.6) |
| CC* (%) | | 100.0 (82.1) | 99.9 (81.8) | 99.6 (79.1) | 99.9 (81.4) |
| **Structure solution and refinement** | | | | | |
| R-work (%) | | 18.61 (52.92) | 22.23 (31.17) | 21.79 (29.59) | 20.69 (35.05) |
| R-free (%) | | 22.08 (58.08) | 26.09 (36.51) | 24.82 (41.06) | 25.83 (37.18) |
| CC-work (%) | | 97.1 (70.5) | 95.3 (75.0) | 90.6 (60.3) | 94.4 (62.0) |
| CC-free (%) | | 96.7 (65.2) | 93.9 (73.7) | 97.4 (54.9) | 92.6 (69.8) |
| R.m.s. deviations | bonds (Å) | 0.030 | 0.006 | 0.029 | 0.003 |
| | angles (°) | 1.05 | 0.66 | 0.52 | 0.53 |
| Average B factor (Å$^2$) | overall | 24.95 | 31.40 | 73.01 | 87.15 |
| | protein | 23.00 | 30.54 | 73.53 | 87.81 |
| | ligand | 39.36 | 45.43 | 56.53 | 65.64 |
| | solvent | 34.24 | 36.13 | N/A | N/A |
| Rotamer outliers (%) | | 0.62 | 0.00 | 0.00 | 0.00 |
| Clashscore | | 0.00 | 1.74 | 0.00 | 0.00 |
| Ramachandran (%) | favored | 97.91 | 98.69 | 99.41 | 98.84 |
| | allowed | 2.09 | 1.31 | 0.59 | 1.16 |
| | outliers | 0.00 | 0.00 | 0.00 | 0.00 |

**Table 1** – Statistics for data collection and processing, structure solution and refinement of the crystal structures of mpox virus poxin and human STING in complexes with MD1203 or MD1202D. Numbers in parentheses refer to the highest resolution shell. R.m.s., root-mean-square.

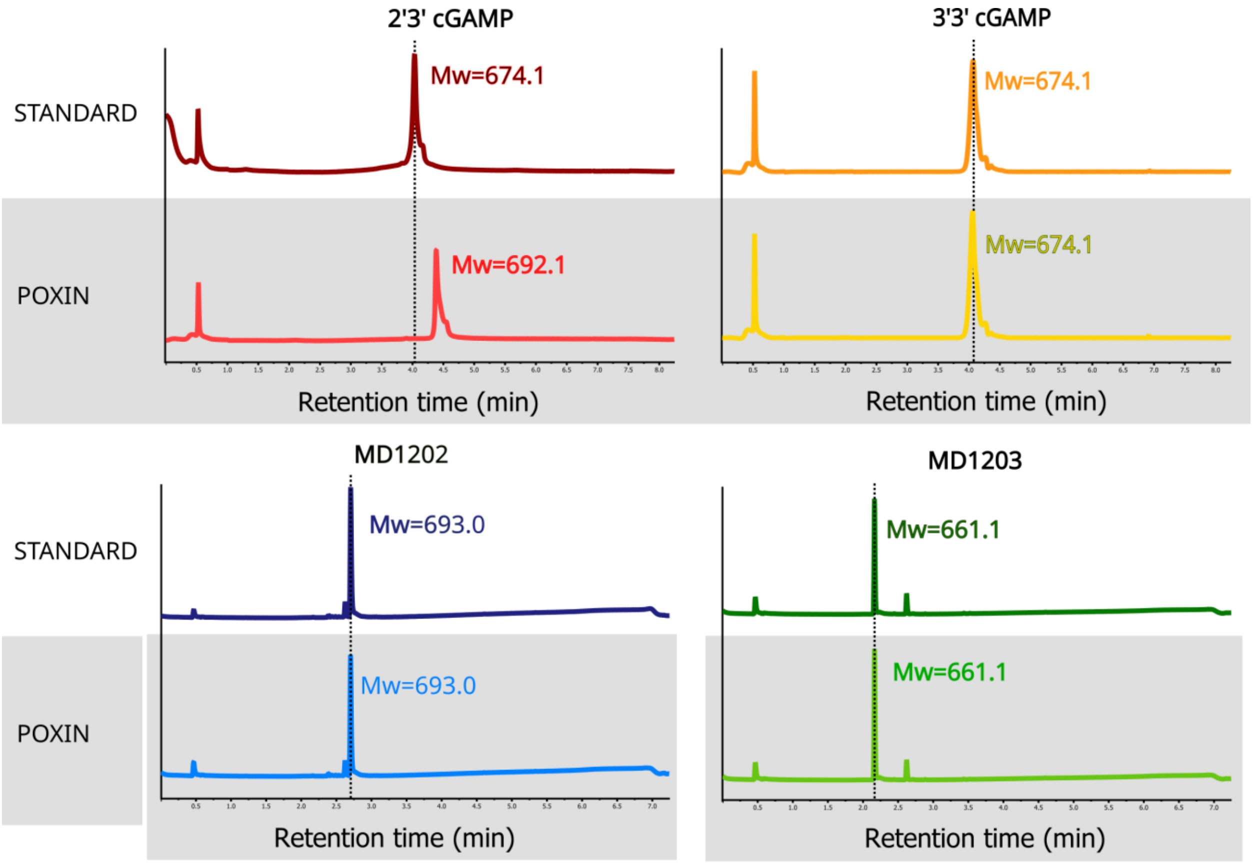


Figure S1. **Poxin does not cleave MD1202D nor MD1203 but cleaves 2',3'-cGAMP**. Each indicated compound was or was not incubated with poxin for one hour and analyzed on LC/MS.

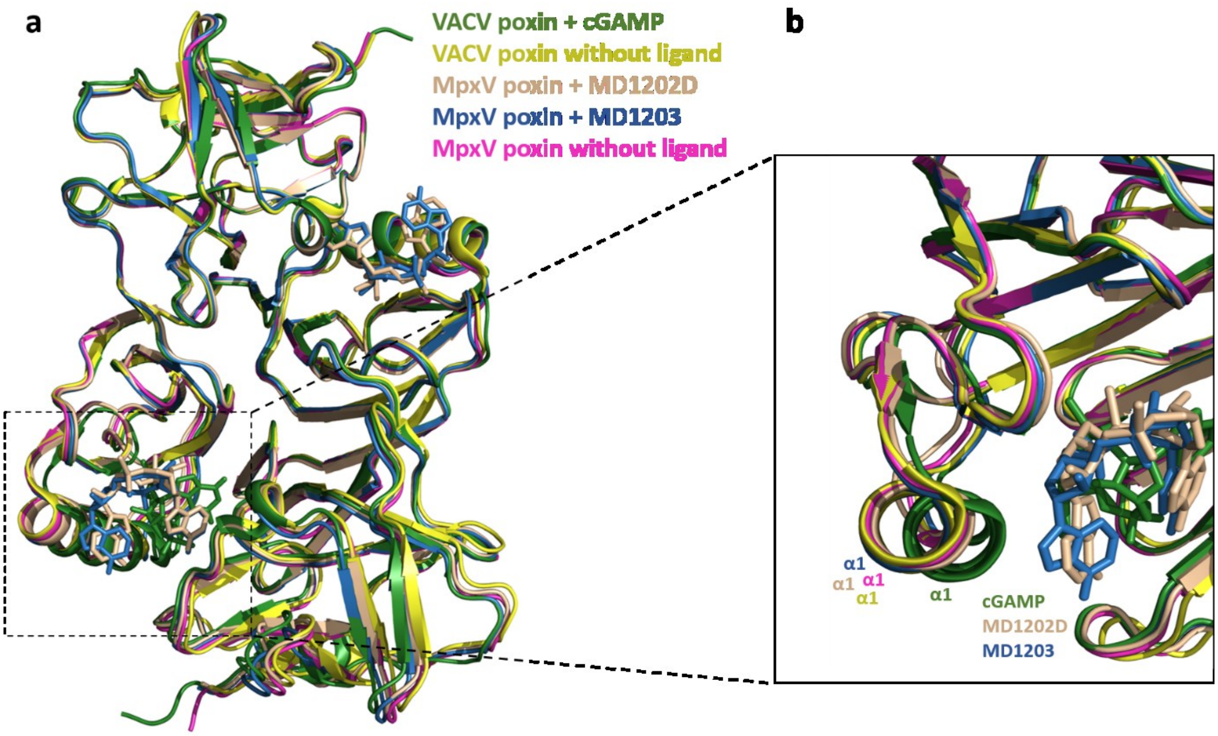


Figure S2. **Conformational changes induced in VACV poxin by ligand binding. a)** Superposition of VACV poxin in complex with cGAMP depicted in green, VACV poxin without ligand depicted in yellow, MpxV poxin in complex with MD1202D depicted in light orange, MpxV poxin in complex with MD1203 depicted in blue and MpxV poxin without ligand depicted in pink. **b)** Close-up on the conformational changes of α-helix 1.
